# Methylphenidate modulates motor cortical dynamics and behavior

**DOI:** 10.1101/2023.10.15.562405

**Authors:** Jessica R. Verhein, Saurabh Vyas, Krishna V. Shenoy

## Abstract

Methylphenidate (MPH, brand: Ritalin) is a common stimulant used both medically and non-medically. Though typically prescribed for its cognitive effects, MPH also affects movement. While it is known that MPH noncompetitively blocks the reuptake of catecholamines through inhibition of dopamine and norepinephrine transporters, a critical step in exploring how it affects behavior is to understand how MPH directly affects neural activity. This would establish an electrophysiological mechanism of action for MPH. Since we now have biologically-grounded network-level hypotheses regarding how populations of motor cortical neurons plan and execute movements, there is a unique opportunity to make testable predictions regarding how systemic MPH administration – a pharmacological perturbation – might affect neural activity in motor cortex. To that end, we administered clinically-relevant doses of MPH to Rhesus monkeys as they performed an instructed-delay reaching task. Concomitantly, we measured neural activity from dorsal premotor and primary motor cortex. Consistent with our predictions, we found dose-dependent and significant effects on reaction time, trial-by-trial variability, and movement speed. We confirmed our hypotheses that changes in reaction time and variability were accompanied by previously established population-level changes in motor cortical preparatory activity and the condition-independent signal that precedes movements. We expected changes in speed to be a result of changes in the amplitude of motor cortical dynamics and/or a translation of those dynamics in activity space. Instead, our data are consistent with a mechanism whereby the neuromodulatory effect of MPH is to increase the gain and/or the signal-to-noise of motor cortical dynamics during reaching. Continued work in this domain to better understand the brain-wide electrophysiological mechanism of action of MPH and other psychoactive drugs could facilitate more targeted treatments for a host of cognitive-motor disorders.

## Introduction

It is estimated that 6.6% of adults in the United States^1^ have used prescription stimulants in a given year, and over 3.5% of children are prescribed stimulants like methylphenidate (MPH; brand name: Ritalin) or d-amphetamine (Adderall) for attention-deficit hyperactivity disorder (ADHD) and related disorders^2^. Many people without clear clinical indications or diagnoses take these drugs illicitly in the hopes of enhancing performance^3–5^. MPH in particular has documented effects on both motor behavior and cognition across multiple mammalian species and has been shown to quicken reaction times (RT)^6–8^, reduce reaction time variability^9,10^, and improve accuracy in behavioral tasks^11,12^. Other stimulants, including caffeine, can decrease motor response times, increase movement velocity, and improve performance in delayed-match-to-sample tasks^13,14^. Methylphenidate has also been shown to reduce RT variability^9,10,15^, improve working memory^16^, quicken RTs associated with making decisions^13^, decrease anticipatory responses^6,9,16^ (but see ref ^17^), and enhance motor steadiness^18^ and grip strength^19^. Recent work also showed an enhancement of implicit, but not explicit, learning on MPH^20^. However, despite many promising routes of investigation and an impressively evolving understanding of the biochemical mechanisms of action of these drugs, it is still largely unclear how and why they can change behavior.

There has been rapid progress over the past decade in uncovering the relationship between cortical population activity and behavior. However, this work has not yet rigorously addressed the effects of cognition-altering medications. While many psychiatric and neurologic medications affect motor behavior and/or cognition, relatively little is understood about the way they affect the activity of single neurons during cognitive tasks. Even less is known about their effects on neural computations in behaving animals^21–24^. Our ultimate goal is to elucidate the effects of methylphenidate (MPH) on motor cortical population activity related to movement preparation and execution. We chose MPH as a behaviorally relevant perturbation likely to affect motor cortical population activity given its documented effects on a broad range of movements across species^6,7,10,19,25,26^, and also because there is clinical and translational value in further understanding the mechanism of action of such a commonly used drug.

The molecular mechanism of action of MPH is well understood. MPH blocks catecholamine reuptake through inhibition of dopamine and norepinephrine transporters^27^. However, its effects on single neurons in awake, behaving monkeys are less well characterized, and its effects on neural population activity during behavior even more poorly so. By comparing neural activity and arm movements with and without systemic administration of MPH, we hope to substantially advance our understanding of the effects of this drug on neural activity and its relationship to behavior. To our knowledge, these experiments constitute one of the first investigations of the mechanism of action of a cognition-altering drug at the level of motor cortical population activity, the first to quantify the effects of MPH on reaching behavior, and the first to assess the effects of MPH on neural population dynamics in behaving monkeys.

In an effort to move beyond molecular mechanisms of MPH, we studied the effects of MPH on the neural computations underlying reaching. In motor cortex in particular, it is not sufficient to correlate single-neuron responses to behavior^28^. This stems largely from the fact that the single-neuron responses rarely match the muscle activity as measured electromyographically (EMG), they vary with movement parameters such as distance and direction in idiosyncratic ways, and they are generally quite heterogeneous^29^. Even upstream, for example in both SMA and M1, single-neuron responses appear to reflect a mixture of task-related variables, suggesting the (erroneous) conclusion that the same general computations are distributed across both regions. Yet careful parcellation of population-level signals reveals that the computations involved are starkly different^30,31^. To tackle such issues, the field of systems neuroscience is embracing a “computation through dynamics” perspective^28^. This framework has pushed the motor field to understand neural activity in terms of population-level signals necessary to reliably generate the desired output. Importantly, these signals tend to be “internal” to the sampled neural population, i.e., they are not directly related to some aspect of the output. Furthermore, these internal signals typically dominate^32^; they may have a larger influence on the response of a typical neuron than the output signals (e.g., force, velocity, etc.) even though the latter are presumably what the network exists to produce.

In dorsal premotor (PMd) and primary motor cortex (M1), the internal signals can be further separated into three types: preparatory, condition-invariant, and execution-related. These signals occupy orthogonal neural dimensions^33,34^, and can thus be isolated by appropriate projections of the population response. Preparatory activity can be interpreted as setting an initial condition to a neural dynamical system; different movements, e.g., leftwards vs. upwards reaches, or slower vs faster reaches, have different initial conditions^35,36,37^. Once initialized, movement is triggered by a precisely timed input from other brain regions, yielding a large translation of the neural state that is ‘condition-invariant’ (i.e., the same regardless of reach type) and a transition to execution-related dynamics^38,39^. Those dynamics display a strong rotational component^40^. This cascade of prepare-trigger-execute applies regardless of how reaching movements are initiated (e.g., self-initiated, external cue-initiated, etc.). While in some cases, preparation can be quite brief, it is obligatory^41^.

Taken together, the field now has a reasonably grounded understanding of the computational role of PMd/M1 during reaching. Indeed, the prepare-trigger-execute series of motor cortical motifs summarized above has been found in a wide range of contexts, including motor learning^42–44^, control of brain-computer interfaces^45^, and speech production^46^. If we could establish that MPH affects reaching behavior in monkeys, then we would be in a particularly well-suited position to understand the causal relationship between MPH administration and motor cortical population-level motifs associated with reaching.

We first established that MPH does indeed affect reaching behavior in Rhesus monkeys. We found that MPH causes dose-dependent and significant effects on trial-by-trial variability, reaction time, and movement speed. Each of these behavioral variables has a direct analog in motor cortical population activity. Thus, we made three neural predictions corresponding to each of these behavioral effects. First, we hypothesized that MPH would directly reduce the variability in population-level preparatory activity. Second, we hypothesized that MPH would reduce the latency of the condition-independent trigger signal. Finally, we hypothesized that MPH would increase the amplitude of movement-period rotational dynamics and/or translate those dynamics to a new location in neural state space.

We confirmed the first two hypotheses using population analyses of the motor cortical responses. Our data were inconsistent with either of our two hypotheses regarding movement speed. That is, previous observations^40,71^ suggested that volitional control of movement speed is associated with a change in the amplitude of rotational dynamics. This follows from the fact that reaching faster isn’t the same as scaling the same neural response to unfold quicker; it requires stronger multi-phasic patterns of muscle activity^36^. Surprisingly, we found that behavioral speed benefits from MPH were accompanied with a change in the frequency of motor cortical dynamics (and not amplitude) and an increase in the signal-to-noise ratio. This suggests that MPH-driven changes in speed likely engage a different mechanism than volitional changes in speed. Our second hypothesis was that MPH may act as a contextual input into motor cortex that causes a shift in neural activity to a new location in state space (presumably to facilitate keeping the dynamics largely unaltered). We found no change in the subspace occupied during preparation or movement under MPH administration, thus rejecting this hypothesis. Instead, our results suggest that the same neural dimensions are engaged during reaching under MPH administration as during the placebo condition, albeit with a change in the dynamics. Taken together, our results are consistent with a mechanism whereby MPH acts as a gain and/or a signal-to-noise modulator on motor cortical neural dynamics.

## Results

We administered oral MPH or placebo in clinically relevant doses (validated with quantitative plasma drug level testing; Table 1) on interleaved days to two adult male Rhesus macaques (monkeys U and P) 15 minutes prior to the start of a center-out delayed reaching task (Figure 1a). The animals were required to withhold reaches to cued targets until presentation of a go cue after a variable delay on each trial and were rewarded with a small bolus of juice at the end of successful trials. In the figures and text that follow, we largely present results for one monkey (U) and one MPH dose (6 mg/kg). The other doses and monkey results are included in Supplementary Materials, and referenced as parentheticals in this section. Note that not all results that follow hold for both animals or both doses. Despite this, our primary findings replicate for both animals, and all results taken together fall along an inferred high-level “inverted-U” shaped dose-response curve, similar to that commonly observed with stimulants. This curve, along with a summary of all results for both monkeys and doses, is plotted in Supplementary Figure 8.

**Figure 1:**
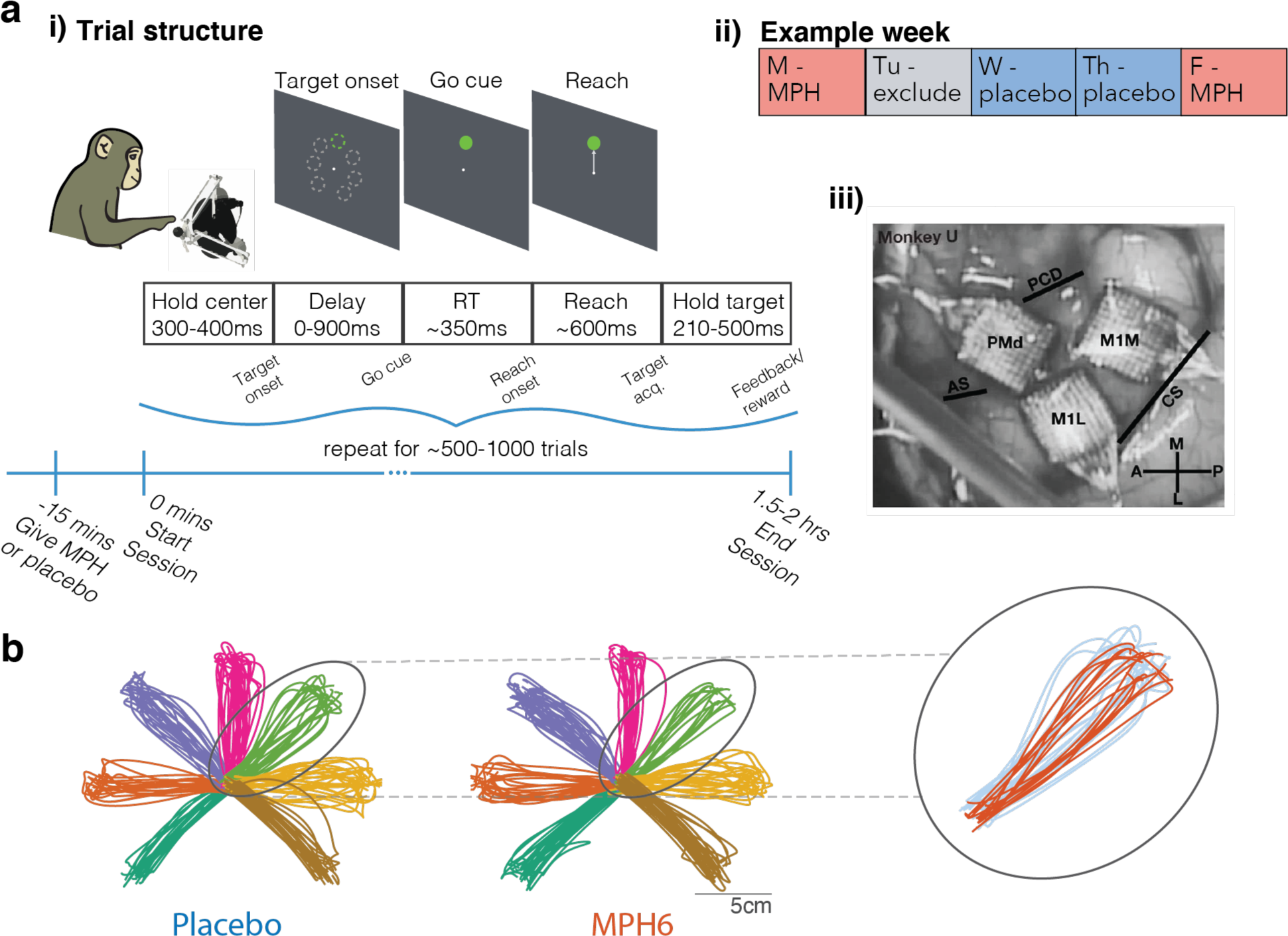
Task and timing. **a) Experimental design. i) Overview of session and trial structure.** Monkeys were given either MPH or placebo 15 min prior to the start of simultaneous behavioral and neural recording sessions. Using a passive manipulandum in a 2D plane with position mapped to a cursor on a screen at eye level, monkeys performed a center-out, delayed reaching task with 7 radial targets. To earn a reward, they were required to withhold reaches to cued targets for a randomized delay period (0ms on 10% of trials; drawn from a uniform 30-900ms distribution on the remaining 90%), during which the reach target jittered on screen. Go cue was represented by cessation of target jittering. Experimenters controlled the session duration to hold trial and reward counts roughly steady across treatments (MPH vs. placebo). **ii) Example session timing.** MPH and placebo sessions were pseudo-randomized to fall on overall similar distributions of weekdays. Sessions the day after MPH sessions were excluded from analysis to minimize potential confounds from stimulant-induced sleep disruption the following day. The experimenter running each session was blinded to the treatment condition. **iii) Chronic electrode array placement.** Photograph from monkey U’s array implantation surgery showing anatomic location of the three 96-channel Utah arrays in PMd and M1, with surrounding cortical surface landmarks (panel reproduced from ^44^). PMd: dorsal premotor cortex; M1m: primary motor cortex, medial array; M1l: primary motor cortex, lateral array; PCD: precentral dimple; AS: arcuate spur; CS: central sulcus. **b) Example reach trajectories by treatment (monkey U, 6 mg/kg dose).** Smoothed 2D hand position trajectories (mapped 1:1 to cursor position on screen) from 180 randomly selected trials per treatment condition. *Left*, placebo sessions; *middle*, MPH sessions. Colors represent different reach targets. *Inset, right:* Example overlaid reach trajectories to a single target from MPH sessions (orange) and placebo sessions (blue).

**Table 1:** Summary of experimental datasets and dosing. Summary of behavioral and electrophysiological datasets collected in each treatment condition (MPH dose and corresponding placebo sessions). Plasma drug levels were obtained separately from behavioral/recording sessions, on a single day in each animal’s home cage, as described in Methods.

| Treatment | # Test sessions | # Control sessions | Trials per session | Plasma MPH level (ng/mL) |  |  |  |
| --- | --- | --- | --- | --- | --- | --- | --- |
|  |  |  |  | @ 30 min | @ 60 min | @ 90 min | @ 110 min |
| Monkey U |  |  |  |  |  |  |  |
| 4.5 mg/kg MPH | 4 | 6 | 539 | n/a | n/a | n/a | n/a |
| 6 mg/kg MPH | 10 | 12 | 525 | 13 | 18.5 | 7.1 | 12.4 |
| Monkey P |  |  |  |  |  |  |  |
| 1.3 mg/kg MPH | 8 | 8 | 1064 | 8.1 | 8.8 | 4.6 | n/a |
| 3 mg/kg MPH | 3 | 5 | 1078 | n/a | n/a | n/a | n/a |

Monkeys were readily able to perform the task in both treatment conditions. Typical reach trajectories to the seven radial targets were grossly unchanged on MPH compared to placebo (see example reaches in Figure 1b), although in aggregate reaches appear slightly less variable on MPH compared to placebo days. Regardless of treatment, reaction times (RT) fell with increasing duration of the instructed delay period (panels ii of Figure 2b and Supplementary Figures 1b, 2b, and 3b), which is evidence that monkeys used the delay period to prepare for the upcoming reach.

**Figure 2:**
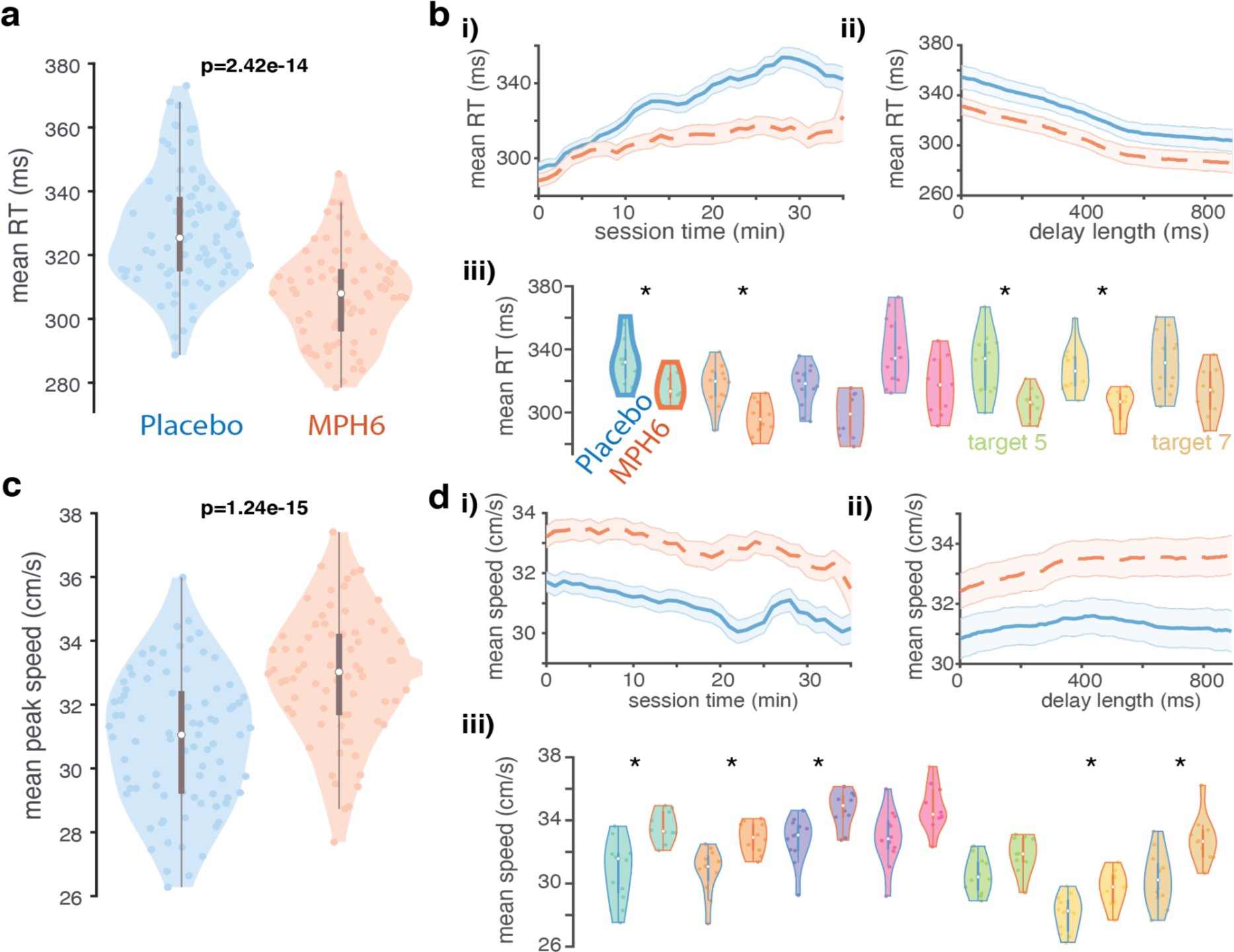
Effects of MPH on reaction times and reach speed (monkey U, 6 mg/kg dose). **a) Reaction time distributions across treatments.** Violin plots showing distributions of mean per-session, per-target RTs in each treatment. Each colored data point is the mean RT across all trials to a single reach target in a single MPH (right, orange) or placebo (left, blue) session. Reaction times were calculated offline as the time from go cue until the hand speed reached 5% of its peak value on a given trial. Box-and-whisker plots for each distribution are overlaid in gray. p-values for differences by treatment condition displayed are found by running ANOVAs on per-session, per-target datapoints including treatment and target as grouping variables, as well as their interaction. **b) RT over time, trial count, delay, and reach target condition across treatments. i) RT over elapsed session time:** moving average of RT across all sessions (orange dashed lines: MPH sessions; blue solid lines: placebo sessions) over increasing per-session elapsed time (in minutes). Session time is truncated to 35min due to limited data in subsequent time windows (secondary to the monkey taking breaks during sessions as well as variability in the overall duration of individual sessions). Error bars: s.e.m. **ii) RT by delay duration:** mean ± s.e.m. of RT calculated over a sliding window across trials with different delay lengths. **iii) RT distributions by reach target.** Conventions for each reach target (pair of violin plots with matching fill colors) as in a, except: 1) p-values were found for differences by treatment in the mean per-session RT values for all trials with a given reach target (rank-sum test). Asterisks denote significance at the .05 level after correction for multiple comparisons. 2) Box-and-whisker plots and violin edges are now color coded by treatment condition (orange for MPH and blue for placebo). **c) Peak reach speed distributions across treatments.** All conventions as in a, except for peak reach speed (also determined on each trial) rather than RT. **d) Peak speed over time, trial count, delay, and reach target condition across treatments.** Same as b, but for peak reach speed.

### MPH causes faster reaction times and reach speeds

We hypothesized that MPH would speed RTs and arm reaches. Consistent with this prediction, we found small but highly significant reductions in RT with MPH compared to placebo sessions (Figure 2a, Supplementary Figures 1a and 2a; but see Supplementary Figure 3a). Peak reach speeds were also significantly faster on MPH (Figure 2c, Supplementary Figures 1c and 2c; but see Supplementary Figure 3c). Both RT and speed effects grossly held up across varying delay lengths (panels ii of Figure 2b,d; Supplementary Figures 1b,d, 2b,d, and 3b,d) and individual reach targets (panels iii of the same figures). While these effect sizes are modest, note that in sports and other related applications, a motor benefit of 5% is highly significant, thus partly contributing to illegal use (e.g., a difference of approximately 5% separated 1^st^ place from 20^th^ place at the 2020 Olympic Games Marathon^47^).

MPH plasma levels rise quickly after oral administration in both humans and monkeys, reaching a peak in monkeys at around 60 minutes^48,49^. We therefore administered MPH (or placebo) 15 minutes prior to the start of each behavioral session, expecting to see some modest initial effect on reach speed and RT that would increase in magnitude over the next ∼45 minutes. RT effects in our data obey these temporal expectations, on average emerging over the first few minutes of the session (panels i of Figure 2b and Supplementary Figures 1b, 2b, and 3b). MPH appears to attenuate a slowing of RT over time and trials seen in placebo sessions. On the other hand, speed effects tend to be present immediately and remain relatively constant from the first few minutes onwards (panels i of Figure 2d and Supplementary Figures 1d and 3d; but see Supplementary Figure 2d). The difference in pharmacodynamics suggests that the effects of the drug on RT and reach speed may have different underlying mechanisms.

Speed and RT effects can be summarized with one statistic, vigor, or the inverse of the time from go cue to target acquisition (RT plus the duration of the reach). Vigor has been shown to correlate with subjective measures of economic value (as measured through choice patterns)^50^. Vigor remained largely stable over sessions within each treatment condition (Supplementary Figures 4a, 1e, 2e, and 3e). This is evidence against tolerance or sensitization developing to MPH, which, if substantial, could dilute or amplify real drug effects in our data, respectively.

### MPH reduces temporal and spatial reach variability

Consistent with prior results from working memory tasks^9,10^, RT variability (quantified as the standard deviation of the RT across all trials to a given reach target within each session) decreased on MPH (Figure 3a, Supplementary Figure 1g; but see Supplementary Figure 2g and 3g), as did peak speed variability (Figure 3b; but see Supplementary Figure 1h, 2h, and 3h). Apart from speed and temporal variability, we hypothesized that MPH would make reach *paths* less variable. To test for this, we focused on two key timepoints (guided by previous work^51,52^) during the reach: the time at which the hand was moving the fastest, and the endpoint. We quantified the spread of the 2D hand position at each of these timepoints across individual trials by using principal components analysis (PCA) to fit an error ellipse capturing on average 90% of the distribution of single-trial hand positions for each target^37,51,52^ (Figure 3c), and then took the area of the resulting ellipses.

**Figure 3:**
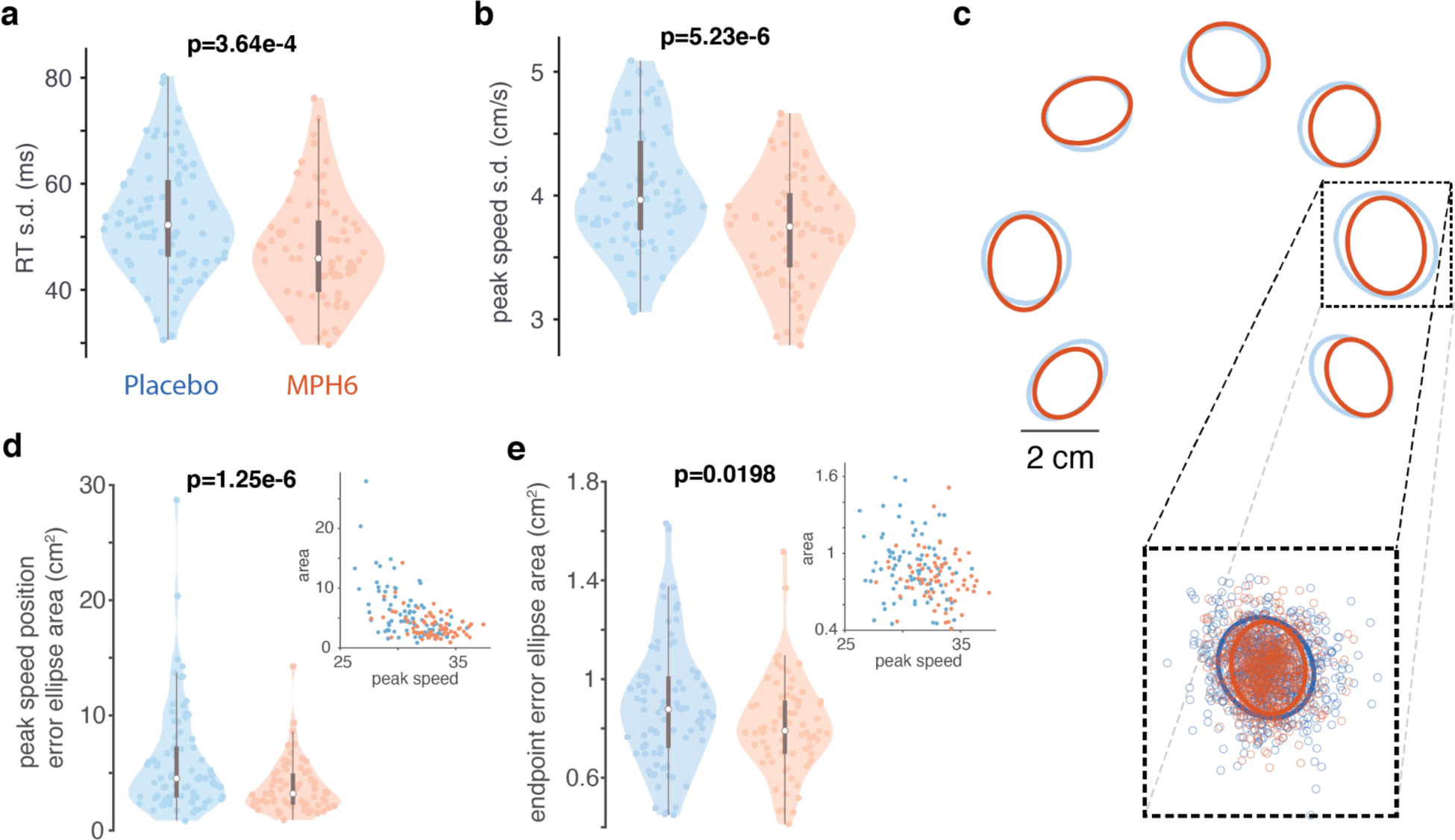
Effects of MPH on reach variability (monkey U, 6 mg/kg dose). **a) RT variability across treatments.** Conventions as in Figure 2a, but for distributions of the per-session, per-target *standard deviation* of the RT in each treatment condition. Each colored data point is the standard deviation of the RT across all trials to a single reach target in a single MPH (right, orange) or placebo (left, blue) session. **b) Peak reach speed variability across treatment conditions.** Same as a, but for distributions of the per-session, per-target standard deviation of the peak reach speed in each treatment. **c-d) Variability of hand position at the time of peak reach speed across treatments. c)** Inset shows full distribution of 2D hand positions at the time of peak reach speed on individual trials in all MPH (orange) and placebo (blue) sessions for an example reach target. Small datapoints represent single-trial hand positions. “Error ellipses (EE)” (in blue for placebo sessions and orange for MPH) found with PCA capture 90% of the variance of the distribution for each target^37,51,52^. **d)** Distributions (by treatment) of the per-session, per-target variability of the hand position at peak speed. Plotting conventions and statistics (ANOVA) as in previous figures. Variability was measured using EE area determined as in a for the distribution of single-trial hand positions for reaches to each target, except on a per-session basis. Each colored data point is the area of the 90% EE fit to the single-trial distributions of hand position at peak speed for all trials to a single reach target in a single MPH session (right, orange), or an associated placebo session (left, blue). Insets: scatter plots of EE area and peak speed (1 data point per session per target; orange: MPH sessions). **e) Variability of reach endpoint position across treatments.** Same as d, but for distributions of 2D hand positions at the endpoint of each reach. Distributions (by treatment) of the per-session, per-target variability of reach endpoint position. Inset: scatter plot of EE area and peak speed (per session per target; orange: MPH).

Computed in this way, hand position variability at the time of peak reach speed was significantly lower on MPH compared to placebo (Figure 3d, Supplementary Figure 1i; but see Supplementary Figure 2i and 3i). Given that there is a well-known speed-accuracy tradeoff for movements^53,54^, and that MPH affected reach speed dose-dependently in our data, we checked for correlations between peak speed and reach variability at peak speed. There was no strong positive correlation in our data (inset scatterplots in Figure 3d; Supplementary Figures 1i, 2i, and 3i). Perhaps unsurprisingly, then, results of the hand-at-peak variability analysis were qualitatively unchanged when crudely adjusted for reach speed (by dividing error ellipse area by peak speed; data not shown). Effects of MPH on endpoint variability were similar to but more modest than effects on hand-at-peak variability, though still significant (Figure 3d-e; but see Supplementary Figures 1j, 2j, and 3j). As above, results were largely unchanged by dividing error ellipse area by peak speed.

### MPH reduces premature movements

Previous studies have reported small effects of MPH enhancing inhibitory control^10,16^. We therefore hypothesized that optimal doses of MPH would reduce premature movements in our task (movements during the enforced delay period, or online detection of RTs under our enforced minimum of 150 ms). Indeed, we found fewer “false starts” on MPH (panels i of Supplementary Figures 4b and 2f; but see Supplementary Figure 1f). At higher doses there were significantly more failed trials resulting from online reaction times that were too *slow* (panels ii of Supplementary Figures 4b, 3f).

### MPH has heterogeneous effects on firing rates

We started by analyzing the effect of MPH on average motor cortical firing rates (FRs). Given the relative stability of recordings from chronically implanted microelectrode arrays across sessions, we were able to compare firing rates of neural activity recorded from the same channels during MPH vs. placebo sessions. While we cannot say for certain that the recordings were from the same populations of individual neurons across days, we have methods to assess that our recorded neural populations were relatively stable (see Methods). Therefore, we were able to compare activity from the same channels under both treatments (MPH vs. placebo). See Figure 4b for some representative example PSTHs calculated across all placebo sessions vs. all MPH sessions. As in these examples, overall task modulation and reach target tuning was generally preserved for most channels across treatments. However, there were often visible differences in baseline FRs and degree of FR modulation by task parameters.

**Figure 4:**
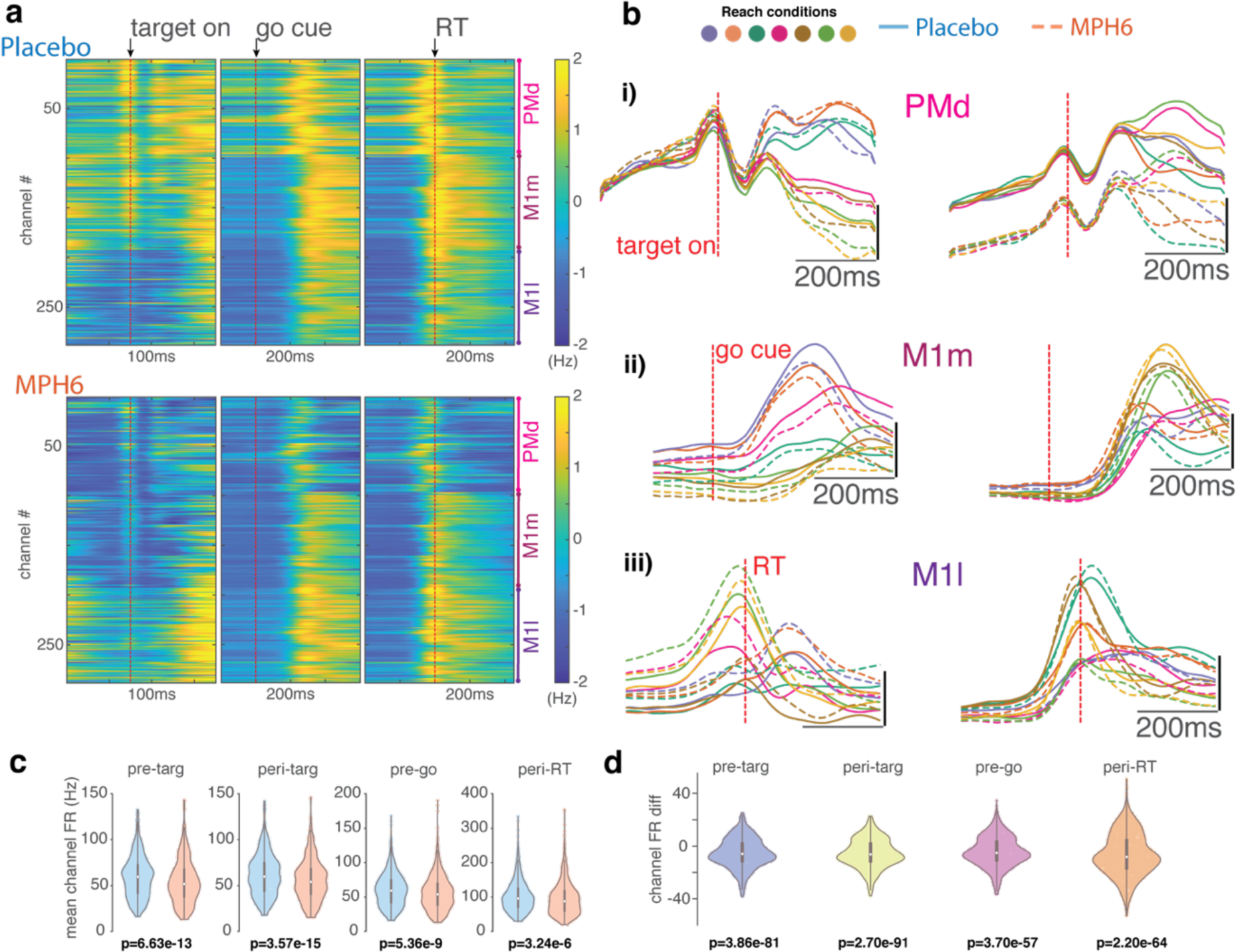
Effects of MPH on motor cortical firing rates (monkey U, 6 mg/kg dose). **a-e) FR differences across treatment conditions. a) FR heatmaps, z-scored across treatments.** Average FRs for each channel (collapsed across all reach targets), z-scored across time and treatment conditions, aligned to key trial epochs around target onset, go cue, and RT). Top, placebo sessions; bottom, MPH. **b) Peri-stimulus time histograms (PSTHs) of representative channels** calculated across all MPH sessions (dashed lines) and all placebo sessions (solid lines). Colors code for the same reach targets as in prior figures. **i)** Two example PMd channels with activity aligned to target onset. **ii)** Go-cue-aligned PSTHs from two M1m channels. **iii)** RT-aligned PSTHs from two M1l channels. **c) Distribution of mean single-channel PSTH values across all time points in each of four trial epochs** (see Methods for exact time windows). One data point per channel per reach target. Box-and-whisker plots overlaid in gray. p-values across all channels were determined using ANOVA with treatment and reach target as grouping variables (and their interaction term). **d) Distribution of single-channel mean condition-averaged FR differences by treatment** across all time points in the epochs from d (difference: mean FR for a single channel and a single reach condition across all trials in MPH sessions minus mean FR from all trials in placebo sessions). Box-and-whisker plots overlaid in gray. P-values here are from Wilcoxon signed rank test.

Consistent with prior work in prefrontal cortex^103^, effects of MPH on motor cortical FRs (unsorted threshold crossings; see ref ^101^) are heterogeneous (Figure 4, Supplementary Figure 5a-d). Overall, we found significantly lower average population FRs on MPH in all trial epochs (Figure 4c-d, Supplementary Figure 5a-d, but see Supplementary Figure 6a-b and 7a-b). This effect may be dependent on dose and exact recording location; in lateral M1 average channel FRs increased on the drug (Figure 4a, Supplementary Figure 5a). We also compared FRs from individual channels during MPH vs. placebo sessions: while the average unit had slightly lower FRs across trial epochs in MPH compared to placebo sessions (a difference of under 10 Hz on average), many channels instead had increased FRs on the drug (Figure 4d, Supplementary Figure 5d).

Having established that MPH causes dose-dependent effects on trial-by-trial variability, reaction time, and movement speed, we made three predictions regarding how MPH may affect motor cortical population activity. That is, we hypothesized that the MPH-driven change in behavior follows from changes in three key dynamical motifs. The three subsections that follow explore these three hypotheses.

### Dynamical Motif I: MPH reduces variability in preparatory activity

We hypothesized that the effect of MPH on trial-by-trial variability would accompany a reduction of variability in the population-level preparatory state in motor cortex. In order to visualize differences in task-related activity with MPH during preparation, we employed dPCA, a dimensionality reduction technique to separate population representations of different task parameters^55^. We first performed dPCA on preparatory activity (smoothed firing rates aligned to target onset) from the entire dataset (concatenated MPH and placebo sessions), including marginalizations by reach target and treatment condition (Figure 5a). Dynamics were largely shared across treatments. There was a large condition-invariant decrease in FRs that temporarily plateaued around target onset, and this neural dimension explained the second-most variance in the data. Reach target population tuning in the top two target-related dimensions (right panels, “Reach condition”) also appeared to emerge slightly faster after target onset on MPH compared to placebo. Note that in this task epoch many of the dPCs were correlated and significantly non-orthogonal (data not shown). This was much less of an issue later in the trial, and perhaps due to the target-aligned activity being lower-dimensional (most variance explained comes from just the top few dimensions).

**Figure 5:**
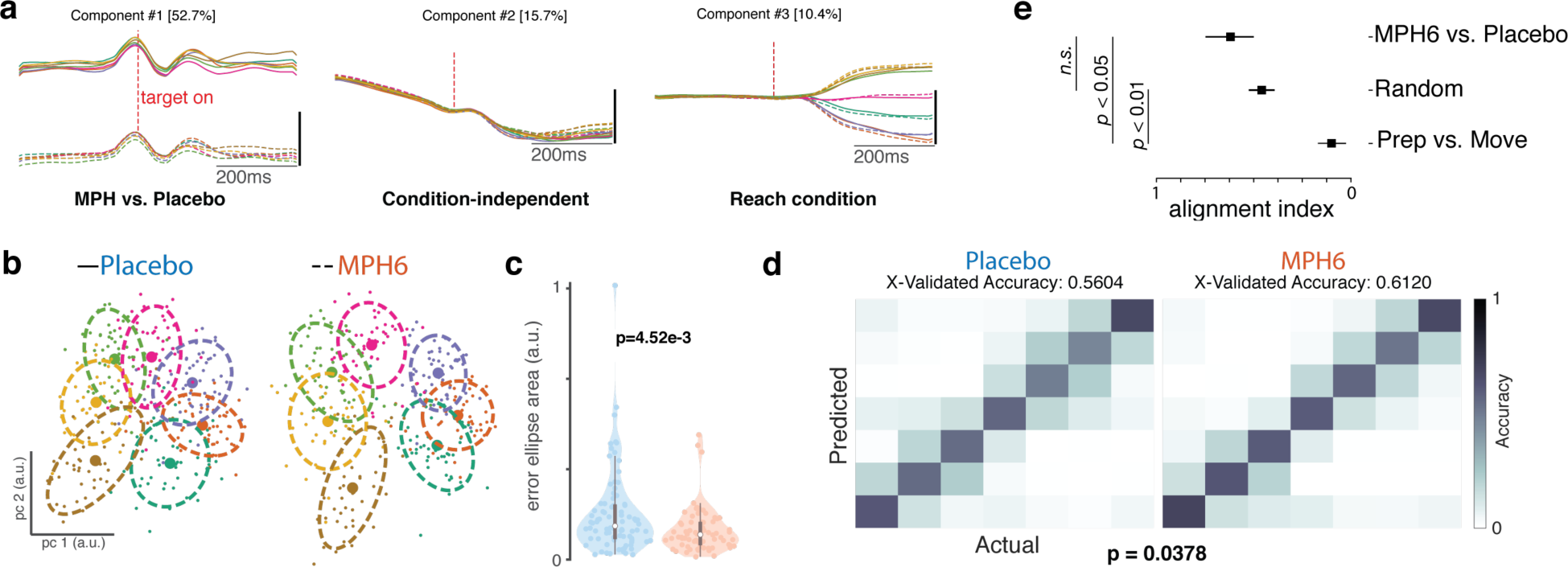
Effects of MPH on motor cortical preparatory activity (monkey U, 6 mg/kg dose). **a) Demixed principal components analysis (dPCA) of U6 data around target onset: top 5 dPCs.** dPCA was performed on smoothed FRs aligned to target onset from a concatenated dataset with all MPH and placebo sessions, for visualization only. dPCA was fit and figure generated using code from ^55^. *Left:* component varying by treatment (MPH vs. placebo); *middle:* condition-independent component; *right:* component varying by reach target. Percentage of variance explained by each dPC is shown in brackets. **b-c) Preparatory activity variability across treatment conditions. b) Example distributions of single-trial preparatory neural states** found using PCA^37,43^ for each reach target (colors) for single sessions in the U6 dataset. *Left:* representative placebo session; *right:* representative MPH session. For each target, small datapoints represent single-trial preparatory neural states in the top two preparatory PCs. Large datapoints represent the centroid, and dashed lines the overlying “error ellipses (EE)” found with PCA and scaled to capture 70% of the variance of the distribution^37^. **c) Distributions (by treatment) of the per-session, per-target variability of the preparatory neural state.** Violin plot plotting conventions and statistics (ANOVA) as in previous figures. Variability was measured using EE area determined as in b) for the distribution of single-trial projections into PC space. Each colored data point is the area of the 70% EE fit to the single-trial PC1-2 projections for all trials to a single reach target in a single MPH session (right, orange) at a given dose, or an associated placebo session (left, blue). **d) Discrete reach target decoding from preparatory activity across treatment conditions.** Gaussian naïve Bayes classification of reach target from raw preparatory activity across all recorded channels. Cross-validated classification^56^ was performed separately for each experimental session. *Left:* classification matrix showing the mean classifier performance across U6 placebo sessions. Colors represent the correlation across trials between the actual target (columns) and the classifier’s predictions (rows). Perfect classification would correspond to ones (yellow) along the diagonal and zeros (dark blue) everywhere else, for a total accuracy of 1. *Right:* Same as left panel, but for MPH sessions. **e) Alignment index.** The alignment index was computed (as described in ^34^) for three different conditions. The bar labeled ‘Prep vs. Move’ corresponds to the preparatory and movement epochs for the MPH data (previously^34^ shown to be orthogonal, i.e., an alignment index value close to zero). The bar labeled ‘Random’ refers to the distribution of indices expected from randomly drawn dimensions from the space occupied by the MPH and placebo data. Finally, the ‘MPH6 vs. Placebo’ bar denotes the alignment index between the preparatory space determined from the MPH and placebo datasets. The error bars for ‘random’ denote 95% confidence interval based on the distribution obtained via bootstrap. The error bars for the other conditions were computed by performing the analysis on a session-by-session basis. To compute the preparatory dimensions, we considered 300ms of time starting 150ms after the onset of the target. The movement dimensions were computed by considering 300ms of time starting 50ms before the onset of the movement, which is an epoch when muscle activity approximately begins to change.

Next, we characterized the variability of the preparatory responses. We performed PCA on 200ms of activity preceding the go cue to find the dimensions in neural state space that explained the most variance for each reach target (see Methods)^37,43^ and then calculated 70% error ellipses of single-trial data projected into the top two PCs (Figure 5b). Error ellipses, calculated per target per session, on average were significantly smaller on MPH (Figure 5c, Supplementary Figure 7c; but see Supplementary Figure 5e and 6c).

We employed a second approach to evaluate treatment effect on population-level preparatory tuning by testing whether the upcoming reach target could be more effectively decoded from the raw delay-period neural activity across all recorded channels on MPH compared to placebo sessions (we would expect so, in the case of broadly enhanced population signal-to-noise). We used a discrete Gaussian naïve Bayes decoder to classify the last 200 ms of delay-period activity for each trial into one of the seven possible reach targets^56^. Target prediction accuracy was several-fold above chance for both treatments, but mean per-session cross-validated accuracy was significantly higher with MPH (Figure 5d; but see Supplementary Figures 5f, 6d and 7d).

Finally, we wanted to test if the preparatory subspace occupied during MPH was the same space that was occupied during the placebo condition. We computed the alignment index^34^, which is a measure of subspace overlap. We started by computing the alignment index between the preparatory and movement epochs. Previous work has demonstrated that these subspaces are orthogonal^34^. We confirmed that this was indeed the case in our data. This also serves as an empirical lower-bound. As in prior work^34^, we performed a second comparison by computing a null distribution of alignment indices. This is done by taking random dimensions within the space occupied by the MPH and placebo data (see ref ^34^). Finally, we computed the alignment index between the MPH and placebo preparatory dimensions; we considered 300ms of time starting 150ms after the onset of the target. We found that the preparatory dimensions occupied during reaches made under placebo are largely similar to those occupied under MPH (Figure 5e).

### Dynamical Motif II: MPH reduces latency of the condition-independent signal

Next, we hypothesized that the effect of MPH on reaction time would accompany changes in the condition-independent signal in motor cortex. We began by visualizing differences in population activity around the time of the prominent “trigger” signal (CIS) as it reliably predicts RT^38^ (Figure 6a). This signal is called conditional-invariant because it reflects movement timing and not identity. We observed a similar CIS across treatments, with separation along the top treatment-related dimension (component #2, bottom panel) largely reflecting baseline FR differences. The top condition-independent component (CIS, top panel) explained the most variance in the data^38^, and rose slightly faster and earlier on MPH. FR differences by treatment were slightly accentuated as CIS rose. Reach-target-dependent components appeared to diverge very slightly faster for different reach targets and returned to baseline faster on MPH.

**Figure 6:**
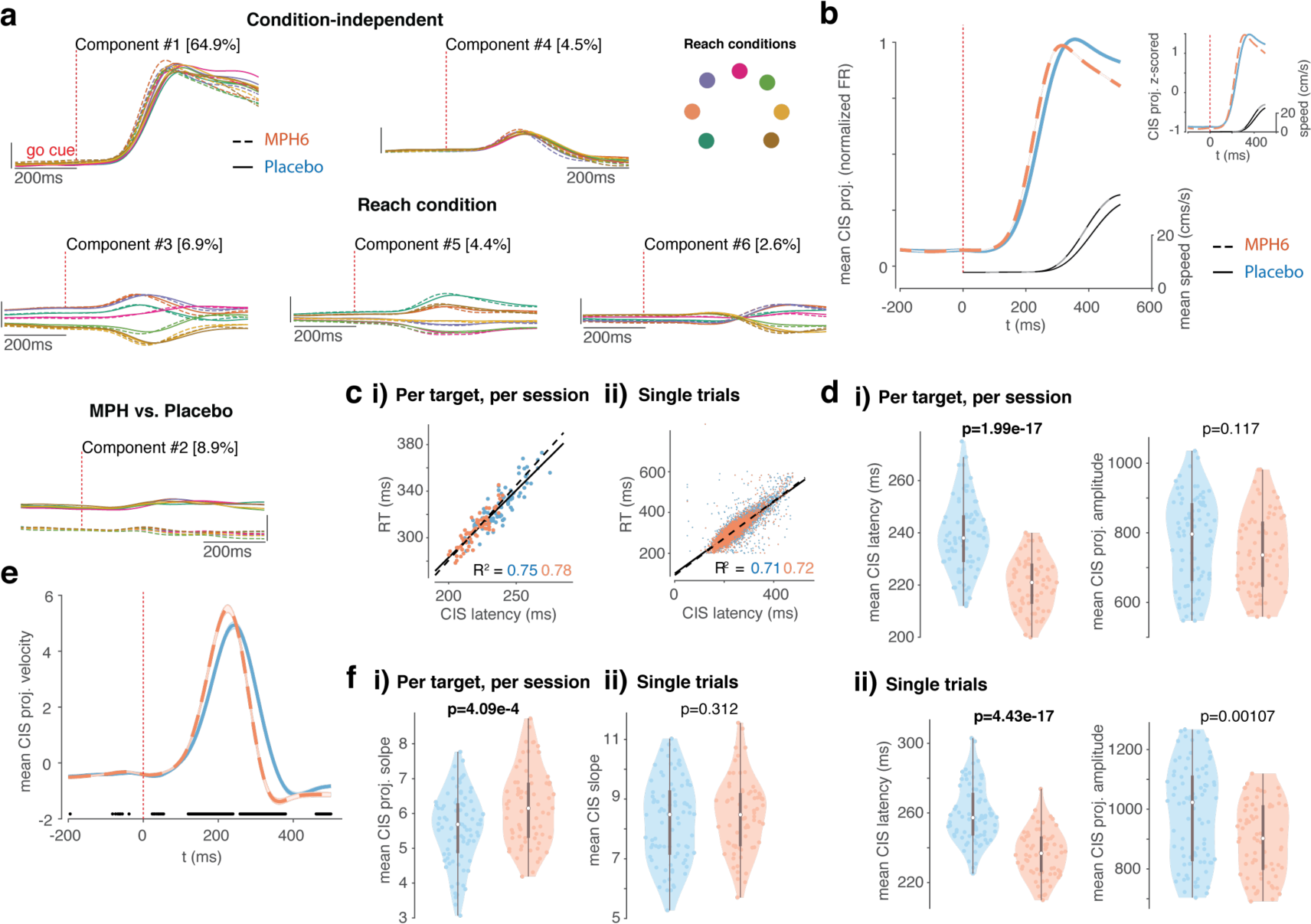
Effects of MPH on motor cortical activity at reach initiation (monkey U, 6 mg/kg dose). **a) Demixed principal components analysis (dPCA) of U6 data around target onset: top 6 dPCs.** Analysis and plotting conventions similar to Figure 5a, except that dPCA was fit on smoothed FRs aligned to go cue. *Top:* condition-independent components; *middle:* components varying by reach target; *bottom:* components varying by treatment. Percentage of variance explained by each dPC is shown in brackets. **b-f) Condition-invariant signal (CIS) across treatment conditions. b)** Mean (± s.e.m.) of projections along the CIS ^38^; the first condition-invariant component, from dPCA fit separately for each treatment) aligned to go cue. Plotting conventions as in Figure 4. Inset in the upper right is from the same data but instead shows the z-scored mean CIS projection for each treatment (normalized amplitude). **c) CIS correlation with reaction time (RT). i)** Correlation between latency of condition-averaged CIS projections and RTs (one data point per session per target). Orange dots and dashed line of best fit correspond to MPH sessions, blue dots and solid line to placebo. **ii)** As in i, but for single trials. d) CIS latency and amplitude across treatments. i) *Left:* Violin plots showing distributions of mean per-session, per-target CIS latencies calculated using condition-averaged trajectories (mean CIS projection per target per session) (the time post-go-cue that the CIS projection reached 50% of its peak on individual trials). Plotting and statistics conventions as in previous figures. *Right:* Same as left, but for CIS projection amplitude. **ii)** Same as i), except for CIS latencies (*left*) and amplitudes (*right*) calculated on a single-trial basis (each data point is the mean single-trial metric across all trials to a given reach target in a given session). **e-f) CIS rate of rise across treatments. e) Mean velocity** (± s.e.m.) of projections along the CIS aligned to go cue, smoothed for visualization only. Plotting conventions as in b. Black dots along the x-axis denote time points at which the MPH and placebo velocity traces significantly differ (false detection rate 0.05). **f) CIS slope across treatments. i)** Violin plots showing distributions of slopes fit to ramping single-session, condition-averaged CIS projections in the 20ms window centered midway to peak (one data point per session per reach target). Plotting and statistics conventions as in d. **ii)** Same as i, but for distributions of mean per-session, per target CIS ramp slopes calculated on single trials.

Similar to prior work^38^, the CIS was found using dPCA, which can separate representation of specific task (and reach) parameters^55^ (see Methods for more details on our approach). Here, dPCA was applied separately on FR tensors from each treatment for each dataset. The CIS is defined as the single condition-independent component that explains the largest amount of variance in the data. We aligned FRs to the RT for increased precision of CIS estimates across conditions, given (1) the CIS relationship with RT is by now well established, and (2) RT and RT variability vary by treatment in our data.

In our data, the CIS appeared to rise slightly earlier and/or faster with MPH (Figure 6b), including after normalizing the amplitude (see z-scored trajectories in the inset). To test for this effect, we first quantified the CIS latency. Single-trial estimates of CIS latency were defined similarly to RT, on a single-trial basis, relative to that trial’s peak CIS amplitude, albeit using a higher threshold (50% of the single-trial CIS projection peak, as opposed to 5% of peak speed for RT) because the single-trial CIS projections were noisier than smoothed reach trajectories. Latencies of condition-averaged CIS projections were defined as on single trials, as the time (from go cue onset) that each trajectory reached 50% of its peak amplitude. Calculated in this way, both single-trial (as in ref ^38^) and condition-averaged CIS latencies were positively correlated with RT (Figure 6c, Supplementary Figure 5h; but see Supplementary Figure 6f and 7f). Both single-trial and condition-averaged CIS latencies were also significantly faster on MPH (left panels of Figure 6d,i-ii, Supplementary Figure 5i,i-ii, and Supplementary Figure 7g; but see Supplementary Figure 6g). Single-trial CIS projection amplitudes were also significantly smaller (right panels of Figure 6d,ii and Supplementary Figure 5i,ii), though once condition-averaged this effect was smaller (right panels of Figure 6d,i and Supplementary Figure 5i,i; but see Supplementary Figure 6g and 7g).

Given that the average CIS trajectory also appeared to rise faster on MPH, we further quantified the speed of this projection. We approached this in two ways: (1) by finding the trajectory velocity in the CIS dimension (taking the derivative of the CIS projection) at each timepoint, and (2) by finding the slope of each trajectory over a longer time window as it rose to a peak (see Methods). The CIS projection velocity, averaged across trials, was indeed significantly faster during the ramp to its peak after go cue onset with MPH, and also decreased earlier post-peak (Figure 6e; but see Supplementary Figure 5j, 6h, and 7h). Despite higher average slopes of the condition-averaged CIS_1_ projections rising to peak with MPH (Figure 6f,i; but see Supplementary Figure 5k,i, 6i, and 7i), the single-trial slopes were not higher (Figure 6f,ii, Supplementary Figure 5k,ii).

### Dynamical Motif III: MPH modulates gain of motor cortical dynamics

Having established that MPH affects reach speed, we hypothesized that there might be interpretable changes to the motor cortical dynamics that accompany reach execution. In particular, we focused on rotational dynamics that are observed during reaching as discovered by Churchland and colleagues^40^. Typically, rotational dynamics are found using a dimensionality reduction method, termed jPCA, where the dynamics matrix is constrained to be skew-symmetric (to capture rotational structure). We used a minor variant of this method in this study (further verified using the HDR algorithm^30^; see Methods). That is, we started by projecting the neural activity around the time of movement onset onto the reach-target-dependent (i.e., “condition-dependent” or CD) dimensions identified using dPCA around the time of the RT (the same procedure used to find the CIS above). Next, we applied jPCA to the condition-averaged CD projections to find rotational structure in the CD dimensions. Significance of results was tested non-parametrically by resampling the data. As in prior work, we focused our analyses to about 200 ms of data starting when preparatory activity transitions to movement-related activity. The results are not sensitive to the exact epochs chosen (data not shown). See Methods for more details.

Our procedure identified rotational dynamics consistent with those previously described – see Figure 7a for trajectories from representative sessions – at frequencies within the range previously reported in M1 (∼8-19 rad/s, or ∼1.5-3Hz)^30,40^. Rotation frequencies were significantly faster on MPH (Figure 7b, Supplementary Figures 5l and 6j; but see Supplementary Figure 7j). Note, for example, that the neural trajectory for each reach condition traverses farther along for MPH (dotted, opaque) than placebo (solid and translucent, Figure 7a).

**Figure 7:**
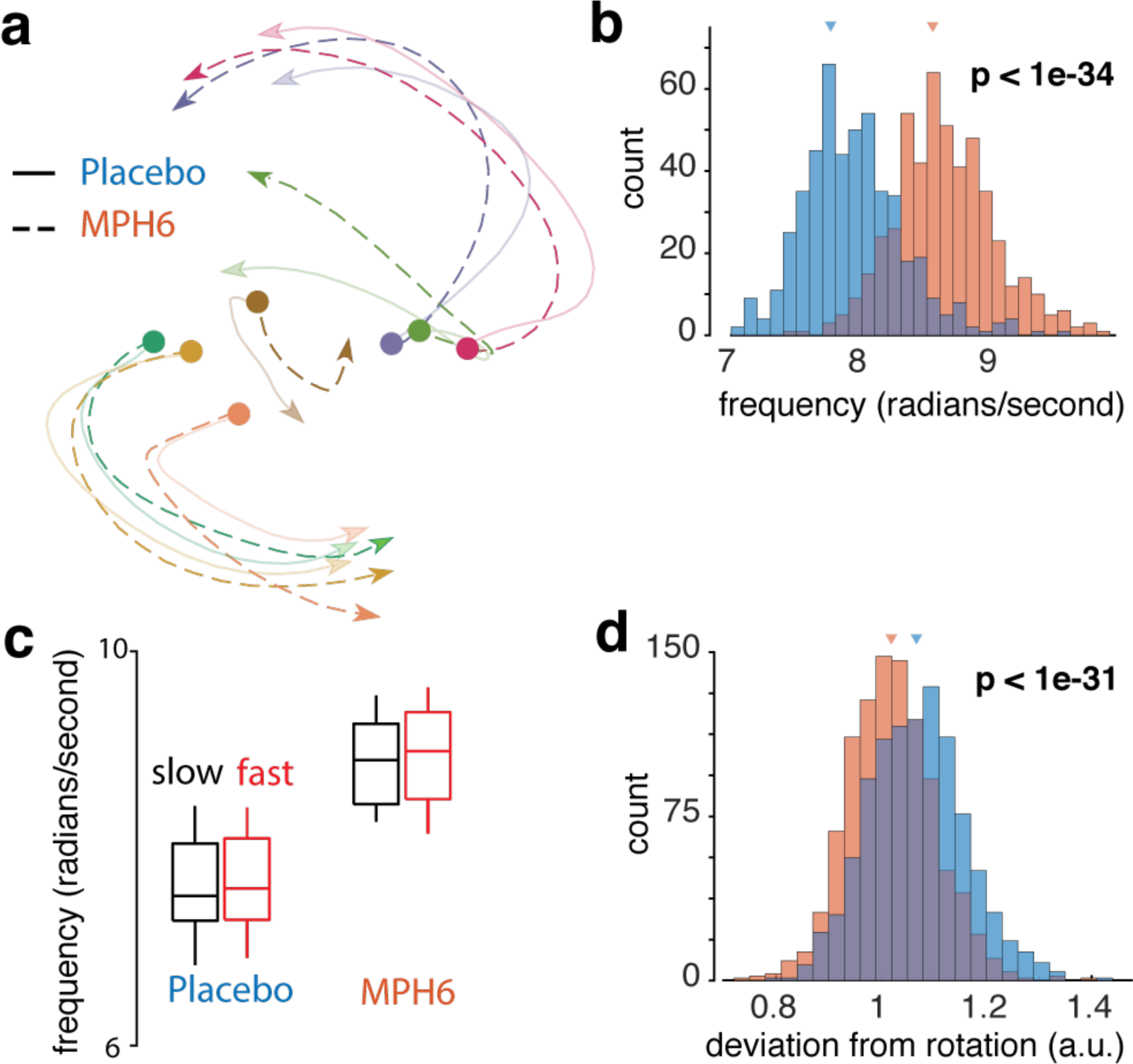
Effects of MPH on rotational dynamics in motor cortex (monkey U, 6 mg/kg dose). Rotational dynamics were found in neural data from 50ms pre- to 150ms post-neural movement onset by applying jPCA to projections into reach-condition-dependent components found using dPCA (see Methods). **a) Example rotational dynamics** from a representative placebo and MPH session. In this example, the initial conditions have been corrected to match across treatment, thus allowing easier comparison of the neural trajectory. Note that the dotted (MPH) trajectories make it farther along the rotation axis in the same amount of time, thus suggesting a higher frequency. Colors represent different reach targets. **b) Rotation frequency across treatments.** Distributions of rotation frequencies calculated from resampled data for each treatment condition. Orange: data from MPH sessions; blue: data from placebo sessions. Triangles mark distribution means. Frequency was calculated from the top 4 dimensions (see Methods). **c) Rotation frequencies are independent of reach speed.** The data for MPH and placebo were split into two groups, each containing either the slowest 10% of trials (black) or the fastest 10% of trials (red). The rotation frequency was computed separately for each group within each treatment. This procedure was repeated across all sessions. **d) Deviation from ideal rotations across treatments.** Deviation (square root of the sum of squared errors) of jPCA rotations from an ideal rotational structure with frequency and amplitude fit to the jPCA trajectories. Bootstrapped deviation distributions from resampled data for each treatment condition. Conventions as in **b)**.

One hypothesis for the apparent increase in rotation frequency could be that this increase is directly caused by increases in reach speed, and critically not a specific result related to MPH. This is unlikely to be the case. A faster reach is not the same as a slower reach with the neural events unfolding at a quicker rate; the movement itself is different and requires generating strong multiphasic patterns of muscle activity (see Churchland et al.^36,40^ for corroborating evidence). Thus, we expected to see a difference in rotation amplitude, and not frequency. Nonetheless, as an additional control, we back-sorted all the placebo sessions into two splits, one group with the slowest 10% of reaches and one group with the fastest 10% of reaches. We fit rotational dynamics to each group separately, and measured any difference in rotation frequency. We found no significant difference. Second, we repeated this analysis for the MPH data, and still found no difference between groups, though the mean for each of the MPH splits was higher than the mean of the placebo splits (Figure 7c).

Based on the dose-dependent effects we observed on reach trajectory variability and neural variability, we tested the hypothesis that MPH might improve the signal-to-noise ratio (SNR) of the rotational dynamics. We define SNR here operationally as the smoothness of the dynamics, where any variability in directions not parallel with the rotation is interpreted as noise. To test this hypothesis, we fit models of idealized circular dynamics to the trajectories (see Methods for details) and calculated the deviation of the measured trajectories from modeled best-fit rotations. Consistent with our hypothesis, the rotations on MPH were closer to modeled best-fit rotations (Figure 7d, Supplementary Figures 5m, 6k and 7k).

## Discussion

The behavioral effects of MPH during reaching afforded a unique opportunity to study its electrophysiological mechanism of action. The computation-through-dynamics framework^28^ has revealed population-level dynamical motifs that are associated with each of the three MPH-induced behavioral effects observed in our study. Thus, we were in a position where we could make testable predictions regarding how MPH may affect motor cortical population activity. In broad strokes, we were able to confirm most of our predictions by analyzing neural activity in two monkeys in a hypothesis-driven manner.

We started by discovering that a low and clinically relevant dose of MPH improves reaction times, increases reach speeds, and reduces reach variability in monkeys performing a delayed reaching task. Together, these results are overall consistent with an “inverted-U” shaped dose-response curve commonly observed with stimulants (see Supplementary Figure 8 and further discussion below for an inferred high-level inverted-U across our animals and doses). Inverted-U dose-response curves have previously been shown to vary depending on the metric of interest, and have often shown substantial inter-individual variability^48,57^, including in monkeys^58^. In keeping with prior evidence pointing to improved inhibitory control with MPH^10,16^, including in monkeys^11^, we also observed some evidence of decreased impulsivity, in the form of fewer premature responses, on the drug.

We observed differences between the RT and speed effects in terms of their temporal evolution during behavioral sessions, with speed effects generally present from the very first trial, but RT effects emerging more slowly over the session. These differences suggest that MPH’s effects on RT and reach speed may have different underlying mechanisms with distinct pharmacodynamics. While this is speculative, in support of this hypothesis, we found that peak speeds on the day after MPH administration (which in and of itself was always a placebo session) still showed statistically significant differences relative to placebo sessions not immediately preceded by MPH sessions (Supplementary Figure 9b). We found no significant changes in most other behavioral variables, including RT (remainder of Supplementary Figure 9).

The RT effects we observed may be tied to improved attention on MPH^24^. It is also possible that the overall vigor-enhancing drug effects we observe could be at least partially tied to MPH-induced changes in reward expectation. Prior studies have shown a connection between movement vigor (including RTs, peak speeds, and amplitude of saccades) and subjective and objective measures of economic value^50,59^. Even the reduced movement variability we saw could be related to reward representations, based on prior work demonstrating that movement variability in healthy volunteers increased with decreased probability of reward^60^. This effect was attenuated in subjects with Parkinson’s disease, suggesting an important role of the basal ganglia in modulating reward-dependent movement variability.

Reductions in movement variability with MPH treatment may also be related to prior reports of decreased behavioral flexibility on MPH^11,61,62^, although there is no cost to making more stereotyped movements in our task. There is evidence in the songbird literature that with less variable song production, such as when reproducing a tutor’s vocalizations, there is increased dopamine in the basal ganglia and decreased variability in basal ganglia outputs^63–65^. Microdialysis and PET imaging studies in monkeys after oral MPH administration have shown significantly increased dopamine levels in the striatum relative to placebo^66,67^; a similar underlying mechanism may thus be at play here. Despite common conceptions that MPH acts selectively on prefrontal cortex, PET imaging in humans using radiolabeled MPH shows by far the highest MPH levels in striatum, and some evidence of MPH binding to dopamine transporters (DAT) in striatum, with diffuse MPH distribution throughout most of the brain (thalamus, cortex, cerebellum), and does not reveal substantially higher MPH levels in prefrontal cortex compared to other cortical regions^68,69^. Other PET work has shown that MPH binds to norepinephrine transporters in the human brain, with highest binding to locus coeruleus, thalamus, and midbrain^70^. Microdialysis and PET imaging studies in monkeys after oral MPH administration have shown significantly increased dopamine levels in the striatum relative to placebo, along with some dose-dependent changes in prefrontal dopamine levels or functional connectivity between the striatum and other cortical and subcortical brain regions^66,67^.

Effects mediated by dopaminergic neurons are certainly not the entire story. MPH blocks reuptake of norepinephrine as well as dopamine, and while norepinephrine receptors are expressed throughout motor cortex, their function in this cortical region is not well understood^74^. At least some behavioral effects of MPH appear to be attributable to modulation of noradrenergic rather than dopaminergic signaling, and norepinephrine has been shown to contribute to inverted-U-shaped dose dependence of SNR modulation in prefrontal cortex^58^. Additionally, even within the better-characterized dopaminergic systems, there appears to be wide variation in distribution of and binding to DAT across different subpopulations of ADHD patients, and MPH’s effects on the dopamine system are very context-dependent^69^.

After establishing that MPH causes behavioral changes in both of our animals in a dose-dependent manner, we made population-level predictions regarding the neural correlates of three behavioral measures: reduced trial-by-trial variability, faster reaction times, and faster reach speeds. Unexpectedly, the two animals and two doses provided distinct but complementary insights. First, in monkey U, preparatory activity variability and CIS latency decreased with MPH administration, while rotational dynamics became faster, with higher signal-to-noise (“SNR,” which we operationally define here as the degree to which the neural trajectory deviated from an idealized rotation about the plane; any off-rotation variability is thus interpreted as “noise”). Second, at the lowest effective dose in monkey P, MPH increased FRs and preparatory activity variability, speed increased along the CIS, and rotational dynamics became modestly faster with increased SNR (Supplementary Figure 6). Intriguingly, at a supratherapeutic dose in monkey P with adverse effects on behavior (Supplementary Figure 3), preparatory variability decreased, CIS latency decreased, and rotations had higher SNR but slower frequency relative to placebo (Supplementary Figure 7).

Our expectations from previous work in computation-through-dynamics^28^ led to a hypothesis that MPH would act as a contextual input that shifts the preparatory and/or movement period dynamics to a new location in neural state space. Indeed, a change in the amplitude of the rotations (as previously observed^40^) would function as such a translation in state space. Our study is perhaps one of the first to report no change in the subspaces occupied during preparation (Figure 5e) or movement (Figure 7), but rather a change in the dynamics themselves. That is, the rotation occurs in the same dimensions, albeit with a different frequency and higher SNR. Taken together across both animals and doses, our results are consistent with a mechanism whereby MPH acts as a gain and/or SNR modulator on behaviorally relevant motor cortical dynamics. Our study cannot go as far as to say that dynamical changes causally drove improvements in the behavior, but we leave this tantalizing hypothesis for future studies to explore.

Our finding of reduced variability during motor preparation on MPH is consistent with reports of cleaner separation of neural trajectories in rat PFC with amphetamine treatment^21^ and overall, with the idea that MPH enhances the SNR of neural activity. The attenuation of the observed CIS amplitude difference by treatment (smaller amplitudes on MPH) for condition-averaged CIS projections could be secondary to more temporal precision in our estimate of the dynamics (e.g., perhaps due in part to the reduced RT variability on MPH) and/or in the spiking activity itself across trials. It is possible that apparent trial-averaged effects on CIS trajectory speed simply reflect more temporally precise estimates of the CIS secondary to decreased variability in the behavior (RT and peak speed) and/or neural activity, rather than truly faster inherent neural dynamics on MPH. This is supported by the finding that, despite higher average slopes of condition-averaged CIS projections rising to peak with MPH, single-trial slopes were not higher.

We also found a clear effect of MPH treatment on rotation frequency that tracks our behavioral finding of increased reach speed. This is in contrast to prior findings that did not find frequency differences by reach speed, but rather increases in rotation amplitude with faster movements^40,71^. The fact that we see a difference in rotation frequency (and not amplitude) is further evidence that MPH acts on the dynamics through a different mechanism than changes driven by volitional state. This might be a fruitful avenue to probe in future experiments. We verified that this difference in frequency was not primarily due to differences in behavior, as opposed to drug effects (Figure 7c). In our dataset the difference between the fastest and slowest reaches is relatively small. In addition, we have a small number of trials when backsorting to the fastest and slowest 10% of trials. This may impact our ability to assess small differences in frequency that may exist in the placebo dataset. For example, it is possible that the fastest and slowest 1% of trials show a difference. While we do not think this is the case (previous studies^40^ found no frequency change even when behavioral differences were larger), future studies should take this into consideration in their experimental design.

We also found an increase in the SNR of the rotational dynamics under MPH. As with the CIS findings, it is quite possible that increased temporal precision of behavior and/or neural activity drive our findings, by allowing for more ready identification of higher-frequency rotational dynamics. Figures 5b and 5d also provide additional evidence for such SNR improvements; the preparatory activity across reach targets was better separated in state space, thus making preparation for upcoming movements more distinguishable. In monkey P, a too-high dose improved SNR of the rotations, while decreasing their frequency (i.e., a dose-dependent decrease in gain). This would suggest that gain and SNR are perhaps not directly yoked.

There are a number of potential circuit-level mechanisms that could accord with our apparent increase in gain. One such possible mechanism, particularly for the condition-independent (reach-nonspecific) effects, could be mediated by MPH’s effects on the ventral tegmental area (VTA). VTA sends dopaminergic inputs, seemingly with glutamatergic co-transmission, to M1^72^. Prior work in rodents found that electrical stimulation of VTA 10ms before otherwise subthreshold stimulation of M1 led to muscle twitches, while stimulating VTA 30ms before otherwise slightly suprathreshold M1 stimulation prevented any EMG response, suggesting that VTA modulates M1 excitability^73^. Previous studies have suggested that this fast excitation-inhibition could increase the SNR in M1 population activity by enhancing temporal spiking precision^74^. In support of this hypothesis, VTA activity is modulated by MPH^75^ as well as by reward expectation^76^. Many of the behavioral effects of MPH we observed, e.g., increased vigor and decreased RT variability, are also seen in cases of increased reward expectation. VTA lesions can ablate at least some behavioral response (increased locomotor activity) to MPH^77^. Activity in VTA, at least in dopaminergic neurons, however, has been largely invariant across different movements and effectors. VTA also projects to striatum, which is consistent with the notion that the basal ganglia may account for at least some of the enhancements in reach execution (such as reduced variability and increased speed) and related dynamics in motor cortex.

### An electrophysiological approach to the mechanism of action

Our study joins recent work in taking important steps toward understanding the mechanism of action of psychotropic drugs, typically approached biochemically, in a relatively novel way: in terms of their effects on neural activity patterns across large neural populations^21–24^. An electrophysiological conception of the mechanism of action is not new to pharmacology: antiarrhythmic and antiepileptic drugs have long been described in this way, and indeed it is broadly accepted that such an approach is essential to understanding how these medications treat pathological electrical activity in the heart and brain, respectively. However, previous approaches to studying therapies for other neurologic, as well as psychiatric, disorders have often not characterized their effects on electrical signaling in the brain, especially at the level of neural population activity. It is clear that at the very least, understanding the effects of psychotropic medications at the tissue, organ, and organ systems levels of organization demands an understanding of their impact on neural activity patterns. It is well established that electrical stimulation of large populations of neurons alone can disrupt behavior and can relieve some neurologic symptoms, such as tremors in Parkinson’s disease or essential tremor, often more effectively than medications. Even nonspecific brain stimulation modalities like electroconvulsive therapy (ECT), which has been around since 1938^78^, can have dramatic effects on behavior. Recently, more targeted modulation of neural activity has shown promising potential for treating refractory psychiatric diseases including depression and obsessive-compulsive disorder (OCD), such as through advances in deep brain stimulation^79–81^. There has also been recent exciting progress toward identifying and targeting electrophysiological biomarkers of psychiatric disease^82,83^ and enhanced approaches to transcranial magnetic stimulation^84^.

A recent study provides a compelling example of the value of this approach^23^. The authors identified an oscillatory electrophysiological signature of the dissociative effects of ketamine, an NMDA-receptor antagonist, in deep layers of the retrosplenial cortex by administering the drug to rodents and observing activity patterns across much of the surface of the brain using widefield imaging. They were then able to identify similar oscillations in a human patient with refractory epilepsy who experienced dissociative seizure auras and reproduce the subjective experience of dissociation with electrical stimulation. Their study shows how understanding the effects of a psychoactive drug at the level of neural activity can enhance our understanding of both the effects of a medical treatment and the covert mental process it modulates across levels of organization.

### Pharmacology as perturbation and known neural effects of stimulants

Other recent studies have also yielded valuable insights into the neural population mechanisms of psychotropic drugs and stimulants. In freely moving rats with low doses of amphetamines, Lapish and colleagues^21^ found evidence of drug-induced separation of neural trajectories in PFC. Hashemnia and colleagues^22^ found that neural trajectories in ACC “contracted” and became less variable. Ni and colleagues^24^ tied MPH-induced improvements in spatial selective attention to reductions in noise correlations in populations of V4 neurons.

Prior to these recent studies, previous work had suggested that MPH might increase cortical firing rates (^85,86^, but not ^87^), increase SNR in cortex^87^, and enhance sensory coding^88^. One previous study of neuronal ensembles in prefrontal cortical responses to CA1/subiculum microstimulation in freely moving rats showed changes in the highest variance dimensions of population activity with systemic MPH administration^86^, providing initial evidence that this drug can affect neural population activity. Electrophysiological data in monkeys to date is limited. However, prior to Ni et al.^24^, one study found diminished representation of reward outcome with systemic MPH treatment in monkeys performing an oculomotor switching task^62^. Another recent study^89^ reported no effect of MPH on prefrontal cortical activity in a visual attention task, either at the single-neuron or population level. It’s worth noting that the doses administered (maximum of 1.3-1.7 mg/kg per monkey) were generally below the range used in other attentional and working memory studies in monkeys. It may also relate to day-to-day variation in behavior, and/or MPH-induced sleep disruption on post-drug days, as each dose was administered on three consecutive days and compared to flanking placebo sessions. A study of atomoxetine, a selective norepinephrine transporter (NET) blocker, in monkeys performing a working memory task found dose-dependent increases in delay-period firing rate for some PFC neurons’ preferred direction^58^ (though this effect could very well be at least partially mediated by effects on dopamine transmission, as the affinity of the NET for dopamine is actually higher than that of the DAT^90^) and blocking NET has been shown to increase prefrontal cortex dopamine levels^91^.

### Limitations and future directions

One limitation of our approach is our relative inability to assess whether our findings represent specific effects of MPH (as opposed to other stimulants) or are specific to motor cortex (as opposed to other connected brain areas). We explicitly set out to test systemic MPH administration and to assess behavioral and neural effects of the drug as it is used in the real world. However, it is important to qualify that the behavioral effects of the drug we observed cannot be specifically or solely attributed to direct effects on motor cortical activity. In fact, there is broad evidence for effects of MPH on many other brain regions, and even an animal’s training history can change the role of different cortical regions in different tasks^92^. We certainly cannot directly attribute any of the signals we identify or perturb to effects of specific cell types or receptors. It is also possible that peripheral effects, including sympathetic arousal^93^, drive some of the changes we observed on MPH. Ideally, future experiments would record from multiple cortical and subcortical brain regions simultaneously to better elucidate network-level dynamics. Future studies should also further compare the motoric effects of MPH with other common stimulating and arousing drugs like caffeine and modafinil, and non-pharmacologically induced arousal states.

We controlled for many non-kinematic but behaviorally relevant variables (such as total reward volume) by ending sessions at a predetermined number of successful trials (except in cases where the monkey refused to continue to work earlier in the session). This prevents us from testing whether MPH affects the amount of time, number of trials, and/or total number of rewards for which our monkeys would continue to work when left to their own devices. Other work has shown that monkeys, when in control of session duration, do perform significantly more trials, particularly at higher doses of MPH^11,24^, consistent with prior work in children with ADHD completing more problems in their schoolwork^57,95,96,97^.

Our task was simple with relatively low demands, especially because our monkeys were very highly trained, with a history of high performance on more complex and difficult reaching tasks (e.g., ref ^43,44^). We did not impose strict endpoint precision criteria, nor any criteria for the precision of reach paths before reaching the targets. It is possible that stronger drug effects on reach trajectory variability might emerge in a task requiring tighter kinematic control. Prior work has shown, for example, both increased variability of movement duration and increased contribution of motor cortical preparatory activity variability to movement duration variability for smaller reach targets^94^. Similarly, while our findings are overall broadly consistent with enhanced motivation and/or attention at optimal MPH doses, more complex, abstract, and cognitively difficult tasks continue to be needed to further elucidate whether MPH has cognitive effects beyond enhancing attention, motivation, and working memory.

Finally, the two doses that monkey P received were either too high (resulting in behavioral deficits within the task – see Supplementary Figure 3 – as well as in the home cage, e.g., reduced appetite), or likely a bit too low (with weak behavioral enhancement; Supplementary Figure 2). More importantly, the recordings in monkey P were of significantly lower quantity and quality compared to monkey U. In monkey U, we recorded from three 96-channel Utah arrays, resulting in 288 channels recorded simultaneously on every session. It’s worth noting that these three arrays are unusually good, even compared to others from our own or other labs in the field. In contrast, in monkey P, we used 16 channel Plexon U/V probes, which resulted in two orders of magnitude fewer channels recorded per session. The number of sessions collected at the too-high dose were limited by the adverse behavioral effects we observed, further reducing the neuron yield. These circumstances made it impossible to perform some of the neural analyses we did for monkey U, particularly the single-trial analyses. Nonetheless, we had enough statistical power to replicate our most critical findings, and thus monkey P represents a replication of our primary behavioral and neural results. Moreover, the too-high dose in monkey P serendipitously provided even greater insight into our proposed mechanisms than pure replication of monkey U’s data would have done.

## Acknowledgements

The authors thank Mark Churchland, Bill Newsome, and Andrew Zimnik for their feedback on this work, Stephen Ryu for help with surgical procedures, Mackenzie Risch and Michelle Wechsler for outstanding veterinary care, and Beverly Davis for administrative assistance. J.R.V. was supported by Stanford MSTP NIH training grant 4T32GM007365, UCSF Psychiatry Research Residency Training Program NIH research education grant R25MH060482, and the Howard Hughes Medical Institute. S.V. was supported by NIH F31 (5F31NS103409) and NIH NINDS F32 (5F32NS124834) grants. K.V.S. was supported through various grants by Howard Hughes Medical Institute, Defense Advanced Research Projects Agency Biological Technology Office, National Institutes of Health, and Simons Foundation Collaboration on the Global Brain.

## Methods

This experiment was performed in two adult male rhesus macaques that had previously been trained on a variety of reaching tasks using a passive manipulandum device (see Figure 1a for task schematic). Data analysis was performed offline (except as required online to run the task as indicated below) using MATLAB v2021b.

### Behavioral Task

The monkeys performed a center-out delayed reaching task (Figure 1a) using a passive manipulandum device (delta.3 haptic device, Force Dimension, Switzerland). Initiation of each trial required holding the device handle (measured by checking for a break in a horizontal light beam directed across the top of the handle) and moving the handle within the 2D experimental plane (45° incline) to direct a cursor on a vertical screen, visible at eye level, to a central fixation dot. The handle was outside of the monkey’s field of view; the device measured hand position and velocity at 1kHz sampling frequency, and cursor movements were coupled to the handle’s movements with 13-20 ms latency. Successful acquisition of the central hold for 400 (300) ms for monkey U (P) was rewarded with a brief juice reward (10-15% of the reward for a successful trial) and triggered the appearance of one of 7 radial reach targets. Targets were evenly spaced at 45° intervals 8 cm from center, with no target at 270° (directly downward). Target presentation was pseudo-randomized in blocks of 7 trials. The initial reach target was a jittering unfilled circle on the screen, serving as a cue during the delay period. Re-acquisition of the central fixation point was permitted if the hand drifted or moved prior to target appearance. To earn a reward, the subject was required to withhold each reach until onset of the “go cue,” in the form of the target ceasing to vacillate and the center filling in. On 10% of trials, there was no delay period (0 ms; go cue was coincident with target appearance and target did not jitter); delays on the remaining 90% of trials were randomly drawn from a uniform distribution between 30-900ms. Online “reaction times” for movement onset were set at subject-specific static speed thresholds. The monkey was given 3 seconds from go cue onset to reach to and acquire the target and hold for 250 (10) ms for monkey U (P) after an enforced 250 (200) ms “settling” period to earn a juice reward. Re-acquisition of the target was permitted within this time frame if the monkey’s hand left the reward zone within a single trial. The trial was aborted if the monkey moved outside of the central fixation target during the delay period, if the online RT was under 150 ms or over 650 (550) ms for monkey U (P), or if the haptic device handle was released at any point throughout the trial. After a brief inter-trial interval of 300 (10) ms for monkey U (P), the monkey was able to return the handle to the center hold position to initiate the next trial.

### Drug administration and dosing

Methylphenidate (brand name: Ritalin) 20 mg immediate release oral tablets were acquired through the Stanford hospital pharmacy. Tablets were cut into halves or quarters as needed to achieve each target dose. Just prior to each experimental session, tablets were ground using a mortar and pestle and the powder was then integrated into a mashed cookie (fig or berry Newtons™) or cookie filling (OREO®). For placebo sessions, the cookie treat was prepared identically except without the MPH powder.

MPH dosing was titrated to behavior for each monkey. Prior work from ^11^ tested a range of doses of oral methylphenidate in three adult macaques in a working memory task and reported significantly higher overall task performance with 3 mg/kg MPH compared to placebo, with a trend toward higher performance at 1.5 mg/kg and significantly impaired performance at 6 mg/kg. Thus, our initial target dosing was 3 mg/kg in each monkey, after testing a single negligible dose (0.3 mg/kg) to monitor for potential adverse reactions. We then monitored for the expected behavioral response (faster RTs and/or reach speeds) to further titrate dosing.

Our two subjects responded differently to the 3 mg/kg dose. Monkey U showed no behavioral response in initial pilot experiments, leading us to titrate up to 4.5 and then 6 mg/kg. Monkey P, on the other hand, started to show adverse effects at 3 mg/kg, including signs of mild irritability and potential anxiety during experimental sessions (widening eyes and quitting experiments early coincident with the expected time of peak drug plasma levels; mild threat posturing when experimenters intervened) and noticeably decreased appetite after returning to the home cage at the end of experiments. These adverse reactions limited our ability to collect a full 3 mg/kg dataset in monkey P as he was unable to reliably complete experimental sessions. Therefore, we titrated down to 1.3 mg/kg for the second dose in this monkey.

### Plasma MPH levels

We sought to validate the resulting dosing with plasma drug levels. We expected to see some variation in dose response across individual subjects, as this is a known source of variability in humans receiving MPH treatment for ADHD and other clinical indications as mentioned above. However, there is significant interindividual variability in GI absorption of MPH in pediatric patients with ADHD^48^, and we suspected that some of the difference between our monkeys may also have been due to differences in absorption of the orally administered drug. In particular, monkey U had a longstanding clinical history of fatty stools, suggestive of malabsorption. Given this suspicion, and given our desire to test clinically relevant MPH doses, we tested plasma levels of the dose that produced the more optimal behavior in each animal (1.3 mg/kg for monkey P and 6 mg/kg for monkey U). Here, MPH was prepared and delivered in mashed cookie, identical to delivery in experimental sessions. The animal was then sedated, blood was drawn at 30, 60, 90, and/or 110 minutes after administration, and blood samples were sent to MedTox laboratories for LabCorp serum methylphenidate level testing via liquid chromatography/tandem mass spectrometry. The resulting plasma levels were compared to a reference range for clinical therapeutic targets in human patients of 5 to ≥20 ng/mL, with potential toxicity starting at 40 ng/mL. Serum levels are not frequently used clinically to dose MPH, with dosing typically titrated to behavior, but can provide a reference for troubleshooting difficulties with dosing or to assess compliance with treatment. However, they have not been convincingly shown to correlate with day-to-day variation in symptom burden for children with ADHD^48^. Results of these serum level tests are presented in Table 1. Except for the 90-min plasma level of the 1.3 mg/kg dose in monkey P, which would be considered subtherapeutic, all measured corresponded to therapeutic serum MPH levels, though the monkey P levels especially were on the low end of the clinically therapeutic range.

### Schedule

The half-life of orally administered MPH in macaques is approximately 107 minutes, with plasma levels peaking around one hour post-administration^49^ and levels in striatum (measured with PET using radiolabeled MPH) peaking around 60-100 min^69^. The prolonged duration of action of this systemically administered pharmacological perturbation precluded counterbalanced experimental blocks of MPH and placebo during individual sessions, as the monkey’s comfort and motivation only permitted restraint for recordings for a few hours at a time (other researchers have faced the same constraints^11,12,24,89,98^). Therefore, MPH and placebo sessions were held on separate days. The conditions were intermixed (rather than, e.g., following back-to-back MPH sessions with back-to-back placebo sessions) to minimize the effects of potential drifts in neural activity and/or behavior over days to weeks, which might have confounded our results. Note that this concern is somewhat validated by differences in the distributions of several of our behavioral metrics, such as RT, in placebo sessions associated with different MPH doses. The monkey was run at the same time of day in both conditions. To control as much as possible for factors like rewards earned/juice consumed and possible performance decrement late in the session (often observed informally and possibly due to development of fatigue over trials), sessions were stopped at a predetermined number of blocks based on the monkey’s natural task engagement over the first several sessions. Post-hoc, sessions were further truncated to the length of the session with the minimum number of completed blocks, across both treatment conditions, to ensure roughly equal trial counts across days for subsequent analyses. Breaks and increases in reward size during behavioral sessions were scheduled to fall within predetermined trial count windows.

Because MPH, like other stimulants, may affect sleep^99^, and sleep deprivation can affect cognitive and motor task performance^100^, we always scheduled one “recovery” day after each MPH session that was not used in either the main MPH or placebo dataset. This also ensured that the monkey never received MPH doses two days in a row, such that well over 20 half-lives elapsed between even the most frequent MPH administrations. Additionally, because in our experience and the anecdotal experience of others, monkeys often perform better or worse on particular days of the week in behavioral experiments (likely associated with facility staffing, experimental workflows, and enrichment schedules), the daily schedule was pseudo-randomized by one of the experimenters to ensure roughly balanced days of the week in each experimental condition.

### Blinding

These experiments were double-blinded with limitations. The monkey received identical-looking treats just prior to all experimental sessions (on placebo, MPH, and recovery days); however, it is possible that he could taste the difference when the treats contained crushed MPH tablets (he was at times slightly more likely to show some resistance to the treat on MPH days, suggesting this was likely the case, at least after completing multiple MPH sessions). Similarly, the lead experimenter running each session was blinded to the daily condition; a different person always made the schedule and prepared the treats. The primary intent of this blinding was to decrease the likelihood of the experimenter running the session or interacting with the monkey differently based on treatment condition. Blinding was preserved through all experimental sessions, and therefore was also preserved for first-pass data analyses. However, it was sometimes possible to guess whether the monkey had received MPH on a given day based on their overall behavior. This was especially the case for the 3 mg/kg dose for monkey P, as it caused him to stop working about an hour into the session, which he did not do at baseline or on non-MPH days.

### Calculation of specific behavioral metrics

Reach kinematics were analyzed post-hoc. Hand position was smoothed using a 4^th^-order lowpass digital Butterworth filter with cutoff frequency of 5 Hz. Speed was calculated as the 2-D derivative of hand position in the x- and y-planes. Reaction time (RT) was defined as the time from go cue onset to 5% of the peak reach speed achieved on a per-trial basis. Trials with RTs less than 200ms or greater than 800ms were excluded from further analysis.

Error ellipses for hand position were found separately for each target as in ^51,52^, as follows. We performed principal components analysis (PCA) on the distribution of 2D cursor position at the time of peak reach speed (or reach endpoint) across all trials to a given target; the first and second principal components (PCs) of each single-target hand position distribution were then scaled back to create axes of ellipses that contained on average 95% of the distribution. Reach curvature/path inefficiency was defined as the path length of the full smoothed reach trajectory on each trial (from target onset through settling on the reach target), divided by the length of a straight line between the center hold point and the reach target (noting that a straight line is not necessarily the most kinetically efficient path). Reach mean squared error (MSE) was calculated as the sum of squared residuals between the single-trial smoothed hand position at each millisecond and a quadratric polynomial curve fit to the mean reach trajectory determined on a per-target, per-treatment basis for the entire dataset (all trials for a given target across days in each treatment group).

RT and peak speed variability were quantified as the standard deviation of the RT and peak speed, respectively, across all trials to a given reach target within an individual session.

### Statistics

Significance of behavioral effects of treatment (MPH vs. placebo) that could be assessed on single trials (RT, peak speed, reach curvature, MSE) was assessed using a separate ANOVA for each metric of interest. Dependent variables were the mean value of the single-trial metric of interest across all trials with a given reach target in each experimental session (one datapoint per reach target per day), using two grouping variables (treatment and reach target) and the interaction between the two. An equivalent approach was used for metrics calculated across trials (error ellipse area, variability of RT and peak speed variability, proportion of trials failed due to premature movement or slow RTs, proportion of successful trials on which the monkey reacquired either the center point or the target), except that the dependent variable for each ANOVA was the single per-target, per-session metric rather than the mean across trials. We adopted this approach rather than comparing full distributions of single-trial metrics to ensure independence of our samples, as the treatment was applied on a per-session rather than a per-trial basis. However, reassuringly, performance for each treatment condition did not differ dramatically across sessions, as in Figure 2g. We used ANOVA with reach target as a separate grouping variable to better isolate the effect of the treatment because most behavioral effects varied substantially by reach target (e.g., RT and peak speed as in Figure 2b and e, panels iv). Indeed, in most cases, we found a significant effect of reach target regardless of the presence or absence of a treatment effect (ANOVA; p-values not shown).

### Electrophysiology

Neural data were recorded using Cerebus recording systems (Blackrock Microsystems, Utah) from three previously and chronically implanted 96-channel Utah arrays (Blackrock) with 1mm electrodes spaced 400µm apart implanted in PMd and medial and lateral M1 (M1m and M1l here, respectively) as in ^44^; see Figure 1a for array placement. Voltage on each channel was sampled at 30kHz and bandpass filtered from 250Hz-7.5kHz, then thresholded at −3.5 times the root-mean-square (RMS).

In monkey P, an acute penetration with a 24-channel linear electrode array (Plexon V-probe) was performed for each session. Recordings were made from left PMd through a small craniotomy (∼7mm AP by 5mm ML) in a previously implanted recording cylinder; see ^43^ for more details regarding cylinder implantation and recording processes. Prior to each behavioral and recording session, a V-probe was slowly advanced, using a NAN micromanipulator, through a non-penetrating blunt guide tube into PMd until neural activity was observed across most channels. The probe was allowed to settle for 45-60 minutes before the session began. The probe was connected to a head stage relaying the voltage to a front-end amplifier (Blackrock), which filtered the signal from 300Hz-7.5kHz, digitized it, and sampled at 30kHz. Each channel was amplified relative to a common reference in the probe. The probe was shorted to the guide tube, and to the metal headpost arm attaching the headpost to the customized primate chair in which the animal was seated, to reduce noise.

### Data preprocessing

As our primary aim was to characterize the effects of MPH on motor cortical population dynamics, and spike sorting has been shown to have little effect on population analyses^101^, no spike sorting was performed. Firing rates for each channel on each trial were smoothed by convolving with a 30-ms standard deviation (s.d.) Gaussian kernel evaluated over 2 s.d.s. For many analyses (dPCA, jPCA, and tangling), firing rates on each channel were “soft normalized” (divided by the range of the channel’s FR plus 5)^40,102^ to partially normalize the dynamic ranges prior to dimensionality reduction but still down-weight contributions from low-FR channels.

For each channel we calculated the mean firing rate (FR) across conditions and time points, the maximal single-reach-target PSTH range, and the maximal single-reach-target s.d. across all trials and time points. A modified SNR was calculated as the FR range divided by the maximal s.d. (adapted from the approach in ^38^). Channels that met three criteria: (1) mean FR of at least 0.5 Hz, (2) FR range of at least 2 Hz, and (3) SNR of at least 0.25 in at least one of three trial epochs (300ms pre- to 400ms post-target onset, 200ms pre- to 600ms post-go cue, or 400ms pre- to 450ms post-RT) were included in further analysis. An additional criterion was imposed: only channels that met the above criteria in *each* treatment condition (e.g., in both MPH and placebo sessions) were included. Still, 100% of channels (288/288) in all monkey U datasets passed screening. In the P1 dataset, 77% (147/192) channels from placebo sessions and 82% (157/192) channels from MPH sessions passed. In the P3 dataset, 84% (101/120) channels from placebo sessions and 85% (61/72) channels from MPH sessions passed.

### Array stability analysis

Prior work in the same monkey, collected just prior to these experiments from the same arrays, showed highly correlated waveforms and preserved directional tuning across 5 sessions in a subset of 71 single units isolated through spike sorting (see Extended Data Fig. 6a-b in ref ^44^). In our dataset, single-channel PSTHs calculated separately from combined drug sessions and combined placebo sessions broadly showed preservation of directional tuning across trial epochs (example PSTHs in Figure 4b are representative). Dimensionality reduction of various forms performed separately on single sessions also showed preservation of tuning at the population level (not shown).

Given that we had strong reason to believe that neural population recordings from monkey U’s chronically implanted Utah arrays were relatively stable across days, we treated them as such (a stable population of the same *N* channels) in many of our neural analyses. Trials collected across separate sessions (different days) were concatenated for each treatment condition, e.g., one firing rate tensor for all placebo sessions (*R_p_* total trials from 12 placebo sessions by *T* timepoints by *N* channels) and another for all sessions with 6mg/kg MPH (*R_m_* total trials from 10 MPH sessions by *T* timepoints by *N* channels), where *T* and *N* are constant across treatments. For monkey P (with acute V-probe recordings from distinct neural populations in each session), no such assumptions could be made.

### Firing rates

To compare firing rate distributions across treatment conditions in different task epochs, we averaged each channel’s smoothed FR across all sessions per treatment condition in the following time windows: ‘pre-targ’ (200 to 100 ms prior to target onset), ‘peri-targ’ (50 ms prior to 50 ms after target onset), ‘pre-go’ (the 200 ms pre-go-cue), and ‘peri-RT’ (50 ms prior to 50 ms after RT). Significance of differences in mean single-channel firing rates was assessed using ANOVA for each time window, with the same grouping variables (treatment and reach target, and the interaction between the two); but dependent variables here for were the mean value of the PSTH for each channel per reach target over the specified time window (one datapoint per channel per reach target).

For z-scored firing rate heat maps (Figure 4a, monkey U only), PSTH tensors for each treatment (*C* reach targets by *T* timepoints by *N* channels) were concatenated, then the full tensor was z-scored across all reach conditions and time points and both treatments.

### Preparatory activity

We used principal components analysis (PCA), an unsupervised dimensionality reduction technique that finds the dimensions in neural state space that explain the most variance in the data, to characterize the preparatory neural population state. We focused on the 200 ms of neural activity just preceding onset of the go cue for this analysis and excluded trials with delays lasting less than 200 ms. For each dataset, as previously described in ^37,43^, firing rates were organized into a *C* × *NT* matrix, where *C* is the number of reach targets (conditions), *N* is the number of neural channels, and *T* is the number of timepoints (here, 200 ms pre-go-cue). We then ran PCA on the firing rate matrix. We plotted the top two PCs (one data point per reach target), then projected single-trial neural activity into the same space (the top two PCs). Error ellipses were found similarly to the process outlined above for hand position distributions and in ^37^ for preparatory neural activity: we performed PCA on the distribution of single-trial projections into PC 1-2 across all trials to a given target; the first and second PCs of each single-target neural state distribution were then scaled back to create axes of ellipses that contained on average 70% of the distribution. Significance of differences in preparatory state variability was assessed with ANOVA as above for reach trajectory variability; dependent variables were the per-target, per-session error ellipse areas.

The alignment index was computed as previously described^34^. Briefly, we grouped all the neural responses into a data matrix Χ ∈ ℝ*^N^*^×*CT*^, where *N* is the total number of neurons, *C* is the total number of conditions, and *T* is the timepoints under consideration. For the preparatory period, we considered *T = 300ms*, starting 150ms after the onset of the target. We computed two such data matrices Χ*_MPH_* and Χ*_placebo_*. We started by performing PCA on the data matrices Χ to obtain a 12-dimensional preparatory subspace. We define a matrix D*_placebo_* as the top 12 prep-PCs obtained under the placebo treatment. We define a matrix C*_MPH_* as the covariance of the matrix X*_MPH_*, where σ*_MPH_*(*i*) is the *i*^th^ singular value of C*_MPH_*. Finally, the alignment index *A* is given by:

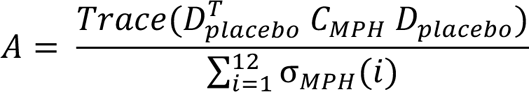

This procedure was repeated where D*_placebo_* was replaced with D*_move_*, where instead of D being comprised of preparatory dimensions from the placebo dataset, it consisted of the top 12 PCs computed by performing PCA on 300ms of movement period data (starting 50ms before the onset of the movement) from the MPH dataset. The ‘random’ data in Figure 5e was computed using a Monte Carlo analysis that sampled random subspaces in the neural state space according to C (which was computed by concatenating the MPH and placebo datasets). The full procedure is described in Supplementary Note 3 of ref ^34^.

### Discrete decoding

We used a Gaussian naïve Bayes classifier to classify vectors of raw firing rates across all recorded neural channels to reach targets, using the general approach from ^56^. We again focused on the last 200ms of preparatory activity, and therefore trials with delay periods under 200ms were again excluded from this analysis. Firing rate vectors consisted of the number of threshold crossings for each channel over the 200ms window prior to go cue onset, divided by the 200ms window width. The log likelihood of observing a given vector of firing rates x under reach target class *C_k_* obeys the following proportion, under assumptions of 1) normally distributed (Gaussian) and statistically independent firing rates across channels and 2) equal variance of each distribution across all *K* classes:

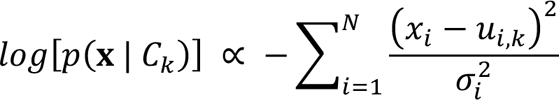

where **x** is the vector of firing rates across *N* channels on a given trial, *i* is the individual channel, *C_k_* is the *k*^th^ class out of *K* = 7 discrete classes (corresponding to reach targets), *u_i,k_* is the mean firing rate for channel *i* in class *k* across all trials in the training data, and 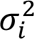 is the variance of the firing rate of channel *i* across all *K* classes. We classified the firing rate vector for each trial to the class (reach target) with the maximum log likelihood after calculating the log likelihood for each class separately. Classification accuracy was determined using ten-fold cross-validation. Classification was performed separately for each session, and significance of accuracy differences across treatment conditions was assessed using a rank-sum test on the vectors of single-session cross-validated accuracies.

### demixed Principal Components Analysis (dPCA)

As above, we used dPCA^55^ to dissect neural population activity into dimensions that reflect different task variables – in particular, time in trial, reach target condition, and in some cases, treatment condition. Briefly, dPCA is a form of dimensionality reduction that captures almost as much variance as PCA but also demixes neural population representations of task parameters. This is achieved in part by decomposing firing rates into averages (“marginalizations”) for related groups of task parameters, and then by using a loss function that penalizes for missed variance in each marginalization as well as in all the other marginalizations and noise. The algorithm is described in more detail in ^55^. We fit dPCA with regularization on all single-trial firing rates; most of our dPCA-based analyses were fit separately on data for each treatment (MPH vs. placebo), and therefore our only marginalizations were for reach target (“condition-dependent,” or CD) and time (“condition-independent”). However, for visualizing differences in population activity across treatments (Figures 5a and 6a, monkey U only), we added marginalizations for treatment and the interaction between treatment and reach target. dPCA was fit with regularization on all single-trial firing rates for monkey U, and separately without regularization mostly on single session PSTHs for monkey P. However, dPCA was run on the full monkey P dataset (PSTHs from all sessions per treatment) to determine the CDs for the analysis of rotational dynamics described below.

### Condition-Independent Signal (CIS)

As described above and similar to ^38^, the CIS was found by performing dPCA separately on smoothed FR tensors from each treatment for each dataset aligned to RT, and then projecting FRs along the top “condition-invariant” dimension (CIS_1_, or just CIS), which is usually the dPCA component that captures the most variance in the dataset. For this analysis, dPCA was fit on data from all trials across days in each treatment condition. All CIS metrics (latency, amplitude, slope, and velocity) were calculated over a time window from 50 to 500ms following go cue onset. Single-trial estimates of CIS latency were found as the time (within that window) at which each projection along CIS1 reached 50% of that trial’s peak CIS amplitude in the same window. Results were comparable if the threshold was decreased, or if it was replaced with 50% of the median CIS peak as in Kaufman et al.^38^ (not shown).

Only condition-averaged CIS projections (for each reach target per session) were analyzed further for monkey P due to excessive noise in single-trial CIS estimates. Latencies of condition-averaged CIS1 projections for both monkeys were defined as on single trials, as the time at which the trajectory reached 50% of its peak amplitude. One MPH session from the P1 dataset (with only 4 channels that passed SNR screening) was excluded from CIS analyses as no condition-invariant components were found with dPCA.

Again, as above, we quantified the CIS projection speed in two ways: (1) by finding the trajectory velocity in the CIS dimension (taking the derivative of the CIS projection) at each timepoint, and (2) by finding the slope of each trajectory (using linear regression on the discrete trajectory timepoints) over a 20 ms window centered around its midway-to-peak point.

### Rotational Dynamics

As described above, we focused our analyses during the “execute” epoch to the previously described rotational dynamics observed in motor cortex. While there is still ongoing work to better understand the nature of these dynamics (e.g., their relationship with sensory and other forms of feedback), they are a sufficiently large signal that arise moments before the onset of movement and persist until the movement is terminated. We adapted the procedure to find this rotational structure originally proposed by Churchland and colleagues^40^.

We analyzed 200ms of neural activity, starting 50ms before the onset of “neural movement” and extending 150ms further. Concretely, the smoothed firing rates on single trials were aligned to the CIS for monkey U, as in^40^, and to the RT for monkey P given his extremely noisy single-trial CIS estimates. The CIS for monkey U was found using dPCA as above. CIS latency here was calculated similarly to the process described in the previous section, except that the more conservative threshold previously reported (halfway to the peak of the median CIS projection) was used for consistency with prior studies^38^ and therefore to increase interpretability of our findings. The smoothed FRs were sampled every 10ms and soft-normalized such that all neurons had similar dynamic range. In contrast to^40^, we did not remove the cross-condition mean and perform PCA. Instead, we projected the data into the “condition-dependent (CD)” (reach-target-dependent) dimensions found using dPCA as above. This yielded a set of nearly independent dimensions that represent different reach targets. We focused our attention to the top 6 (P1, P3, and U6 datasets) or the top 8 (U4 dataset) CD dimensions. CD dimensions 7-8 were added for U4 as, in each treatment condition in this dataset, there was a much higher-frequency rotational component across CDs 6-8, each of which captured a relatively small proportion of the variance in the data. This component was present in CD 6 for the MPH data but in CD 7-8 for the placebo data, and therefore greatly skewed our results toward much higher rotational frequencies with MPH if we only considered CD 1-6. We condition-averaged the data to yield an average firing rate tensor (conditions x time x neurons). For each condition separately, we projected the trial-averaged data onto the subspace spanned by the top 6 (or 8) dPCA-derived dimensions. The resulting data - which represents a 6 (or 8) dimensional neural state for every time and condition - was then fed into the jPCA algorithm^40^. The result of jPCA is a dynamics matrix that has been constrained to be skew symmetric. This guarantees that the resulting eigenvalues are purely imaginary and thus this captures rotational structure in the data – here, in the CD dimensions. We analyzed the eigenvalues and computed the corresponding rotation frequency.

In order to create a distribution of rotation frequencies, we followed a simple resampling procedure. For all sessions, we randomly sampled (with replacement) 20% of the trials. This was done separately for the placebo sessions and the MPH sessions. Using the resulting trials, we fit a jPCA model and computed the rotation frequency. We performed 1000 such random samplings and generated a distribution for MPH frequencies and placebo frequencies. We verified that these distributions were normal before applying a *t*-test. Similar results were found when placebo and MPH sessions were mixed to compute the null distribution.

To assess the degree to which the neural trajectories deviated from an ideal rotational structure, we fit a rotational model where the amplitude and phase were a function of initial condition. We made the simplifying assumption that under an ideal rotational model the neural trajectory would make a perfect circle once it departed from the jPC1 dimension. Note that in the jPCA procedure, the initial conditions splay out across jPC1 and then rotate counter-clockwise (by design). We assumed that once the trajectory was non-zero in jPC2, it would rotate counter-clockwise and make a perfect circle. We computed the deviation (defined as the square root of the sum of the squared difference) between this ideal rotation and the actual path that the neural trajectory took. The hypothesis we tested was whether neural trajectories following MPH had a smaller rotational deviation than placebo. Note that the ideal rotational model was fit to minimize the difference between an ideal circular trajectory and the actual trajectory, where the frequency and amplitude for the ideal trajectory were chosen to minimize the deviation from the actual trajectory. We performed a standard bootstrap procedure with replacement (1000 samplings) and generated a histogram of deviations (across all reach conditions). After checking for normality, we performed a *t*-test.

We repeated many of our analyses using the Hypothesis-guided Dimensionality Reduction (HDR) method described by Lara and colleagues^30^. The HDR method neither requires removing the cross-condition mean nor requires an explicit skew-symmetric constraint, and thus does not explicitly look for rotations. Furthermore, it jointly estimates the dynamical and condition-invariant dimensions. We were able to replicate all of our findings using this more general method, which we interpret as a sanity check (data not shown).

---

**Supplementary Figure 1:**
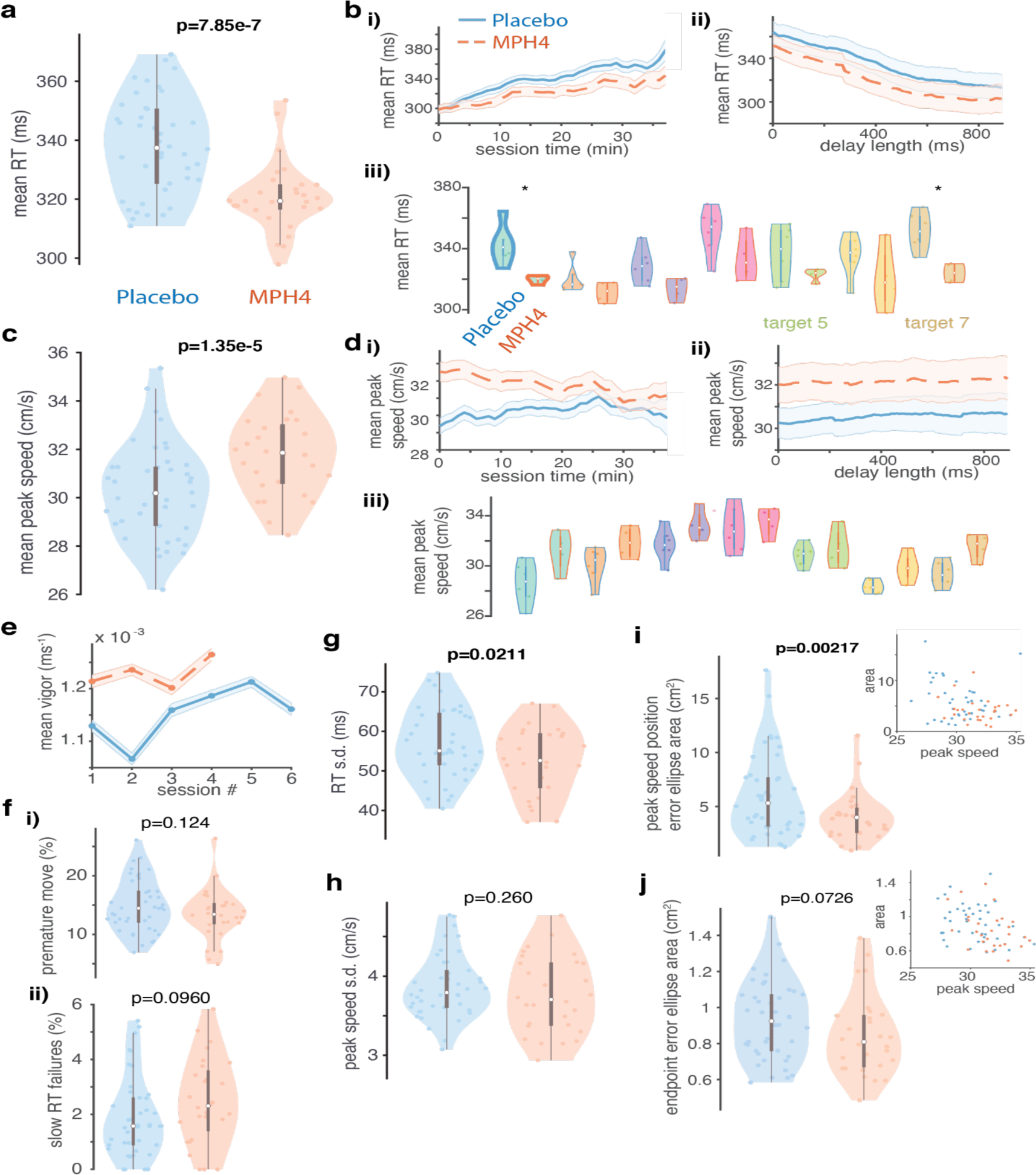
Behavioral effects of MPH for the 4 mg/kg MPH dose in monkey U and associated placebo sessions (“U4” dataset). **a) Reaction time distributions across treatments.** Same as Figure 2a, but for U4. **b) RT over elapsed session time, delay, and reach target condition across treatments.** Same as Figure 2b, but for U4. **c) Peak reach speed distributions across treatments.** Same as Figure 2c, but for U4. **d) Peak speed over time, trial count, delay, and reach target condition across treatments.** Same as b, but for peak reach speed. **e) Vigor over sessions by treatment.** Mean per-session vigor (± s.e.m.) across all trials for each treatment. Vigor is calculated per trial as the inverse of the sum of the RT and the reach duration ^50,59^. Orange dashed lines: MPH sessions; blue solid lines: placebo sessions. **f) Impulsivity. i) False starts across treatments.** Conventions as in Figure 2a, but for distributions of the per-session, per-target proportion of trials aborted due to premature hand movements (moving during the delay period, or online RT <150ms) in each treatment. **ii) Sluggish starts across treatments.** Same as i, but for per-session, per-target proportion of trials aborted due to online RTs that were too slow. **g) RT variability across treatments.** Same as Figure 3a, but for U4. **h) Peak reach speed variability across treatment conditions.** Same as g, but for distributions of the per-session, per-target standard deviation of the peak reach speed in each treatment. **i) Variability of hand position at the time of peak reach speed across treatments.** Same as Figure 3d, but for U4. **j) Variability of reach endpoint position across treatments.** Same as i, but for distributions of 2D hand positions at the endpoint of each reach.

**Supplementary Figure 2:**
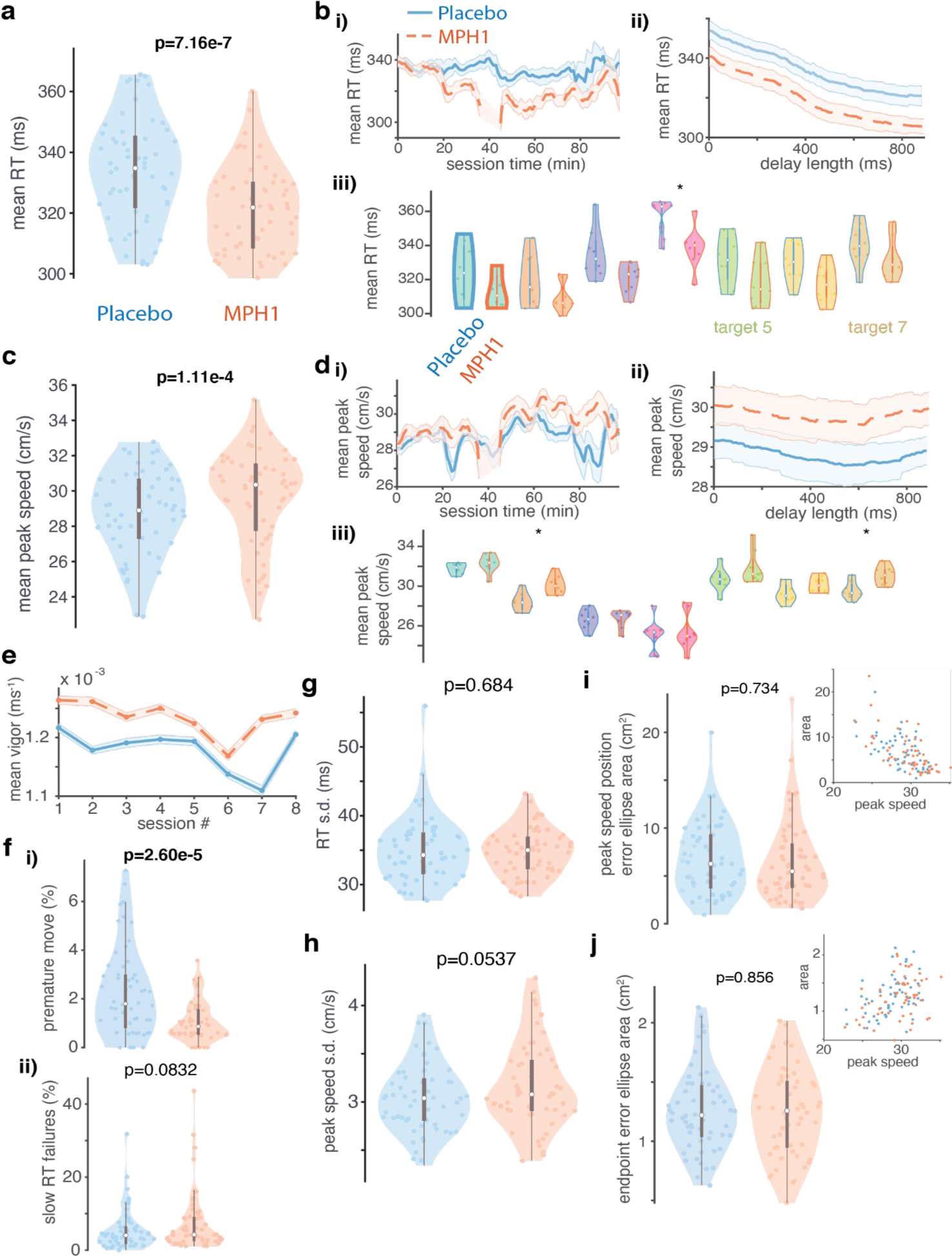
Behavioral effects of MPH for the 1.3 mg/kg MPH dose in monkey P and associated placebo sessions (“P1” dataset). **a) Reaction time distributions across treatments.** Same as Figure 2a, but for P1. b) RT over elapsed session time, delay, and reach target condition across treatments. Same as Figure 2b, but for P1. c) Peak reach speed distributions across treatments. Same as Figure 2c, but for P1. d) Peak speed over time, trial count, delay, and reach target condition across treatments. Same as b, but for peak reach speed. e) Vigor over sessions by treatment. Same as Supplementary Figure 1e, but for P3. f) Impulsivity. **i) False starts across treatments.** Same as Supplementary Figure 1f, but for P3. **ii) Sluggish starts across treatments.** Same as i, but for per-session, per-target proportion of trials aborted due to online RTs that were too slow. **g) RT variability across treatments.** Same as Figure 3a, but for P1. **h) Peak reach speed variability across treatment conditions.** Same as g, but for distributions of the per-session, per-target standard deviation of the peak reach speed in each treatment. **i) Variability of hand position at the time of peak reach speed across treatments.** Same as Figure 3d, but for P1. **j) Variability of reach endpoint position across treatments.** Same as i, but for distributions of 2D hand positions at the endpoint of each reach.

**Supplementary Figure 3:**
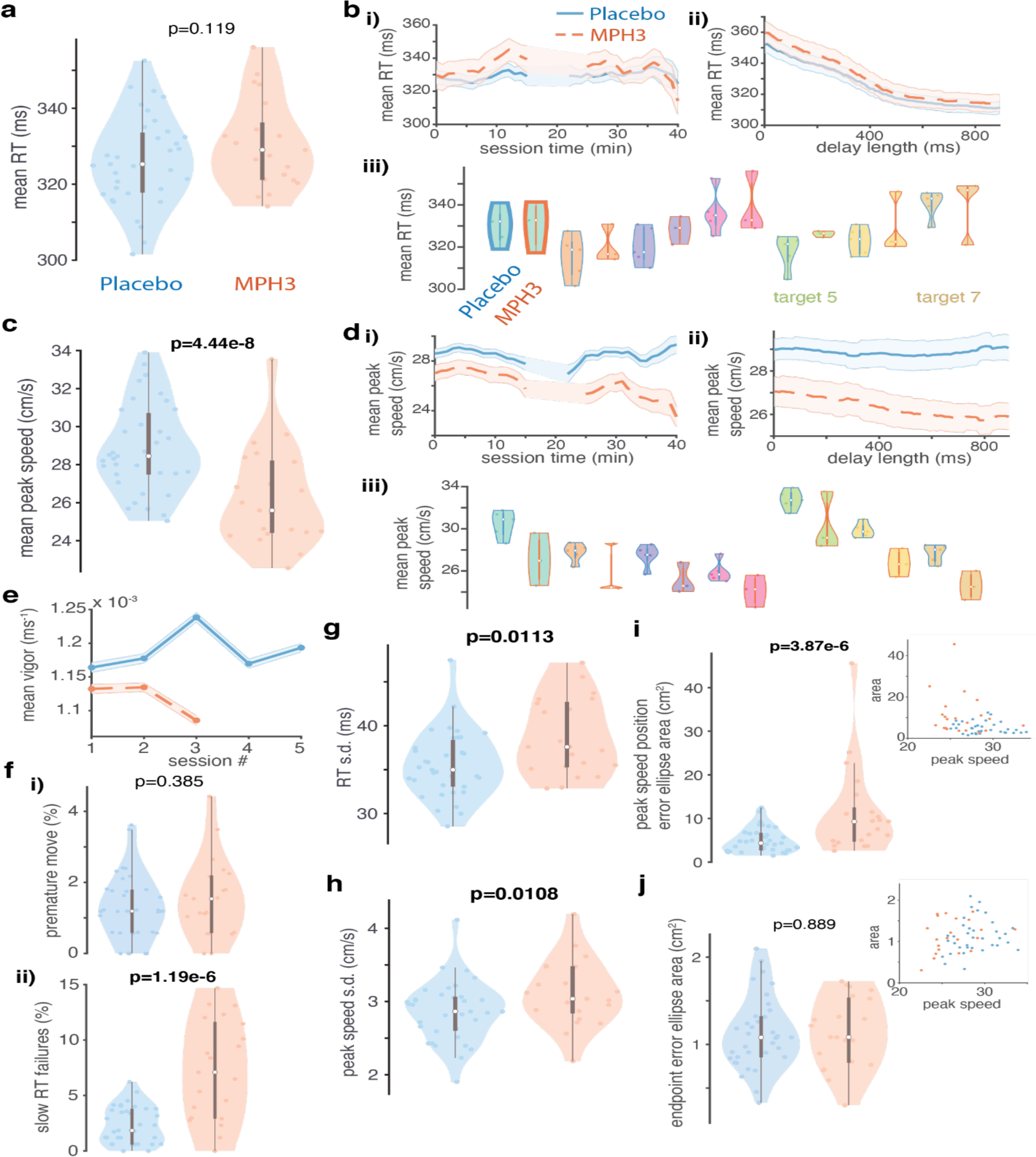
Behavioral effects of MPH for the 3 mg/kg MPH dose in monkey P and associated placebo sessions (“P3” dataset). **a) Reaction time distributions across treatments.** Same as Figure 2a, but for P3. **b) RT over elapsed session time, delay, and reach target condition across treatments.** Same as Figure 2b, but for P3. **c) Peak reach speed distributions across treatments.** Same as Figure 2c, but for P3. **d) Peak speed over time, trial count, delay, and reach target condition across treatments.** Same as b, but for peak reach speed. **e) Vigor over sessions by treatment.** Same as Supplementary Figure 1e, but for P3. **f) Impulsivity. i) False starts across treatments.** Same as Supplementary Figure 1f, but for P3. **ii) Sluggish starts across treatments.** Same as i, but for per-session, per-target proportion of trials aborted due to online RTs that were too slow. **g) RT variability across treatments.** Same as Figure 3a, but for P3. **h) Peak reach speed variability across treatment conditions.** Same as g, but for distributions of the per-session, per-target standard deviation of the peak reach speed in each treatment. **i) Variability of hand position at the time of peak reach speed across treatments.** Same as Figure 3d, but for P3. **j) Variability of reach endpoint position across treatments.** Same as i, but for distributions of 2D hand positions at the endpoint of each reach.

**Supplementary Figure 4:**
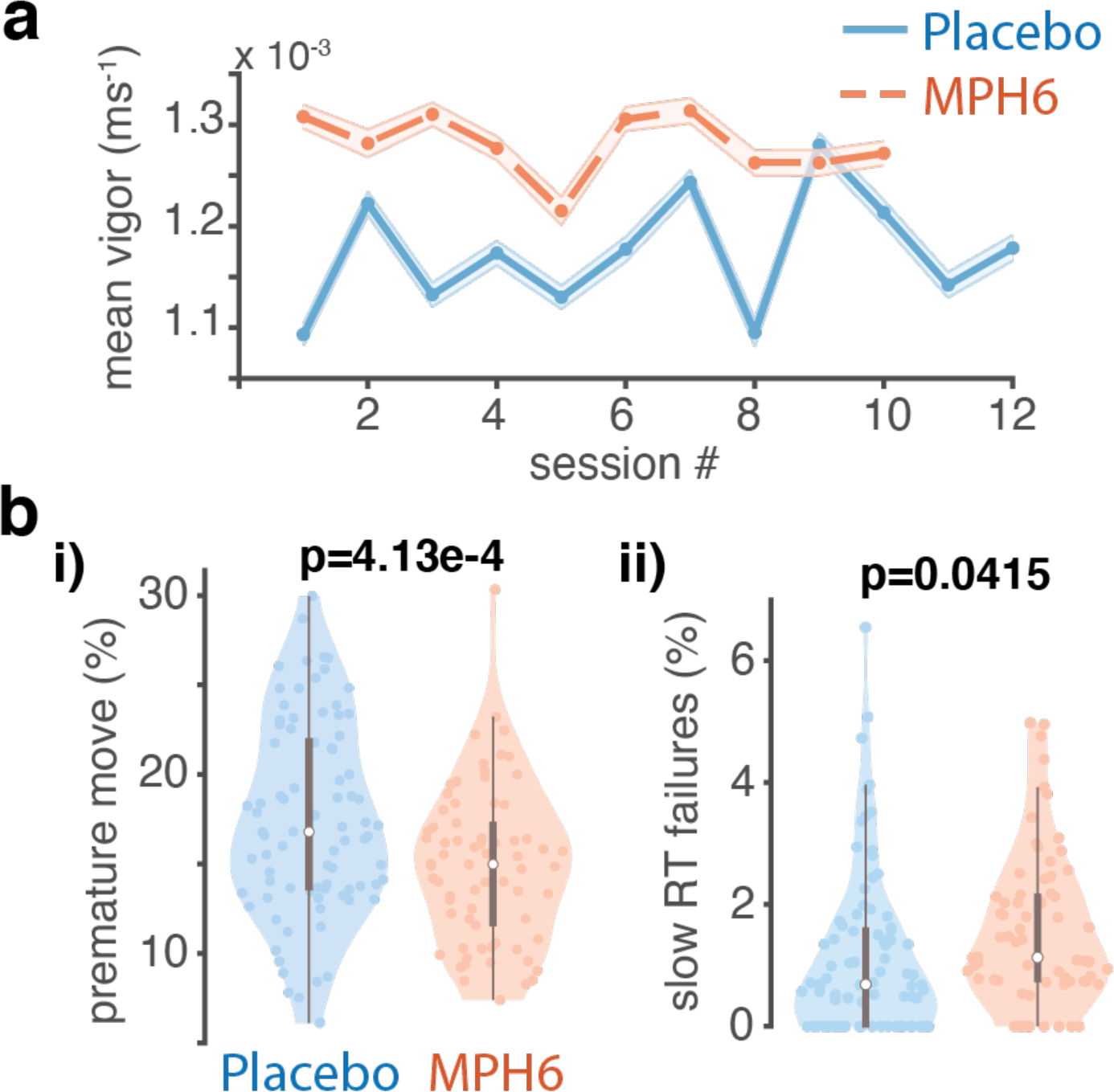
Tolerance and trial outcomes. **a) Vigor over sessions by treatment.** Mean per-session vigor (± s.e.m.) across all trials for each treatment. Vigor is calculated per trial as the inverse of the sum of the RT and the reach duration ^50,59^. Orange dashed lines: MPH sessions; blue solid lines: placebo sessions. **b) Impulsivity. i) False starts across treatments.** Conventions as in Figure 2a, but for distributions of the per-session, per-target proportion of trials aborted due to premature hand movements (moving during the delay period, or online RT <150ms) in each treatment. **ii) Sluggish starts across treatments.** Same as i, but for per-session, per-target proportion of trials aborted due to online RTs that were too slow.

**Supplementary Figure 5:**
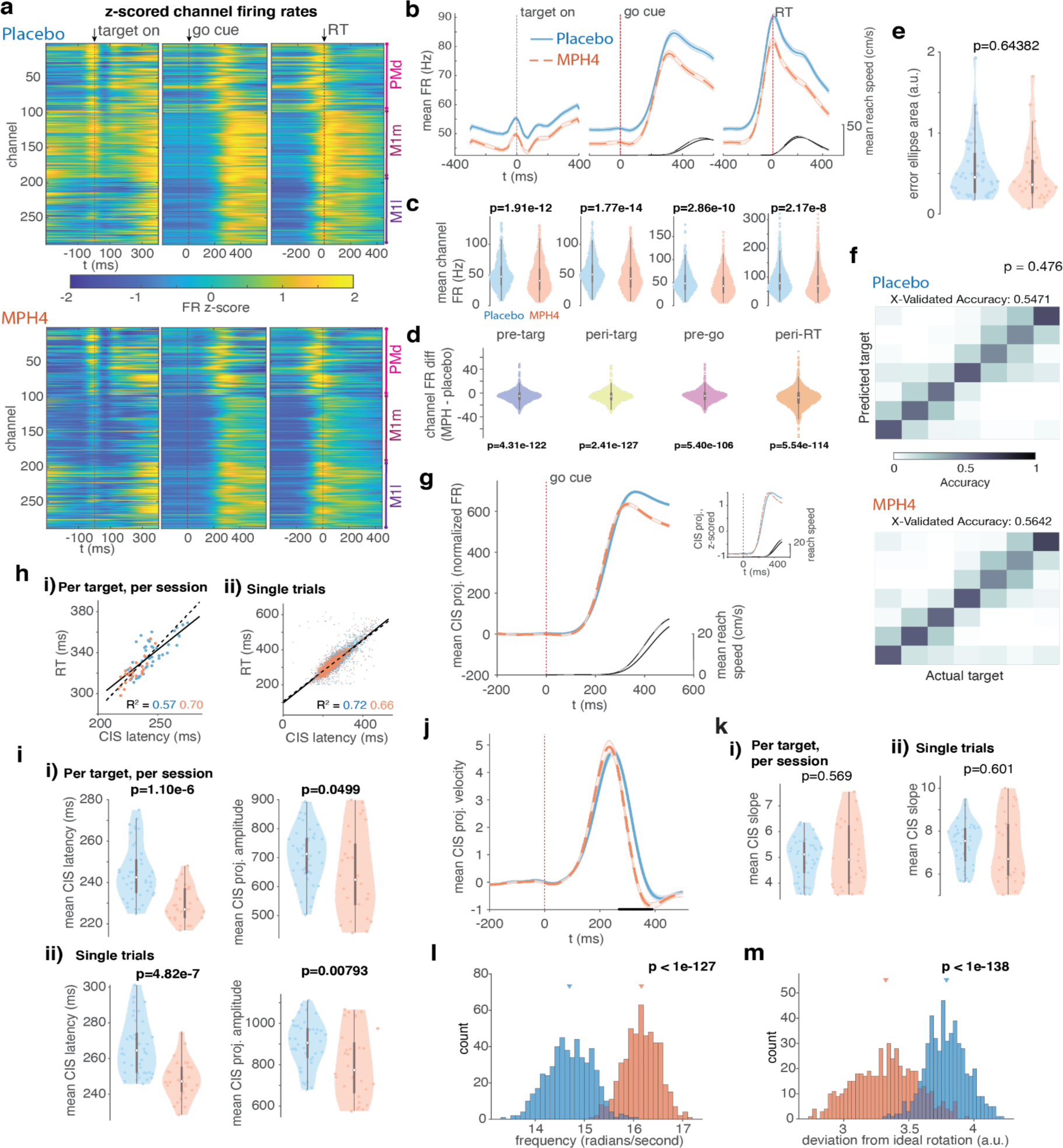
Neural effects of MPH for the U4 dataset. **a-e) FR differences across treatment conditions. a) FR heatmaps, z-scored across treatments.** Same as Figure 4a, but for U4. **b) Mean population FR trajectories by treatment.** Collapsed across all reach targets and aligned to target onset, go cue, and RT, across all channels during MPH vs. placebo sessions. Mean reach speed trajectories are overlaid on the right axes. Dashed lines: MPH; solid lines: placebo sessions. **c) Distribution of mean single-channel PSTH values across all time points in each of four trial epochs.** Same as Figure 4d, but for U4. **d) Distribution of single-channel mean condition-averaged FR differences by treatment.** Same as Figure 4d, but for U4. **e) Distributions (by treatment) of the per-session, per-target variability of the preparatory neural state.** Same as Figure 5c, but for U4. **f) Discrete reach target decoding from preparatory activity across treatment conditions.** Same as Figure 5d, but for U4. **g) Condition-invariant signal (CIS) across treatment conditions.** Same as Figure 6b, but for U4. **h) CIS correlation with reaction time (RT).** Same as Figure 6c, but for U4. **i) CIS latency and amplitude across treatments.** Same as Figure 6d, but for U4. **j) CIS velocity across treatments.** Same as Figure 6e, but for U4. **k) CIS slope across treatments.** Same as Figure 6f, but for U4. **l) Execution-related rotation frequency across treatments.** Same as Figure 7b, but for U4, with one exception: frequency was calculated from the top 6 dimensions (see Methods). **m) Deviation from ideal rotations across treatments.** Same as Figure 7c, but for U4, and with deviation calculated from the top 6 dimensions as in l.

**Supplementary Figure 6:**
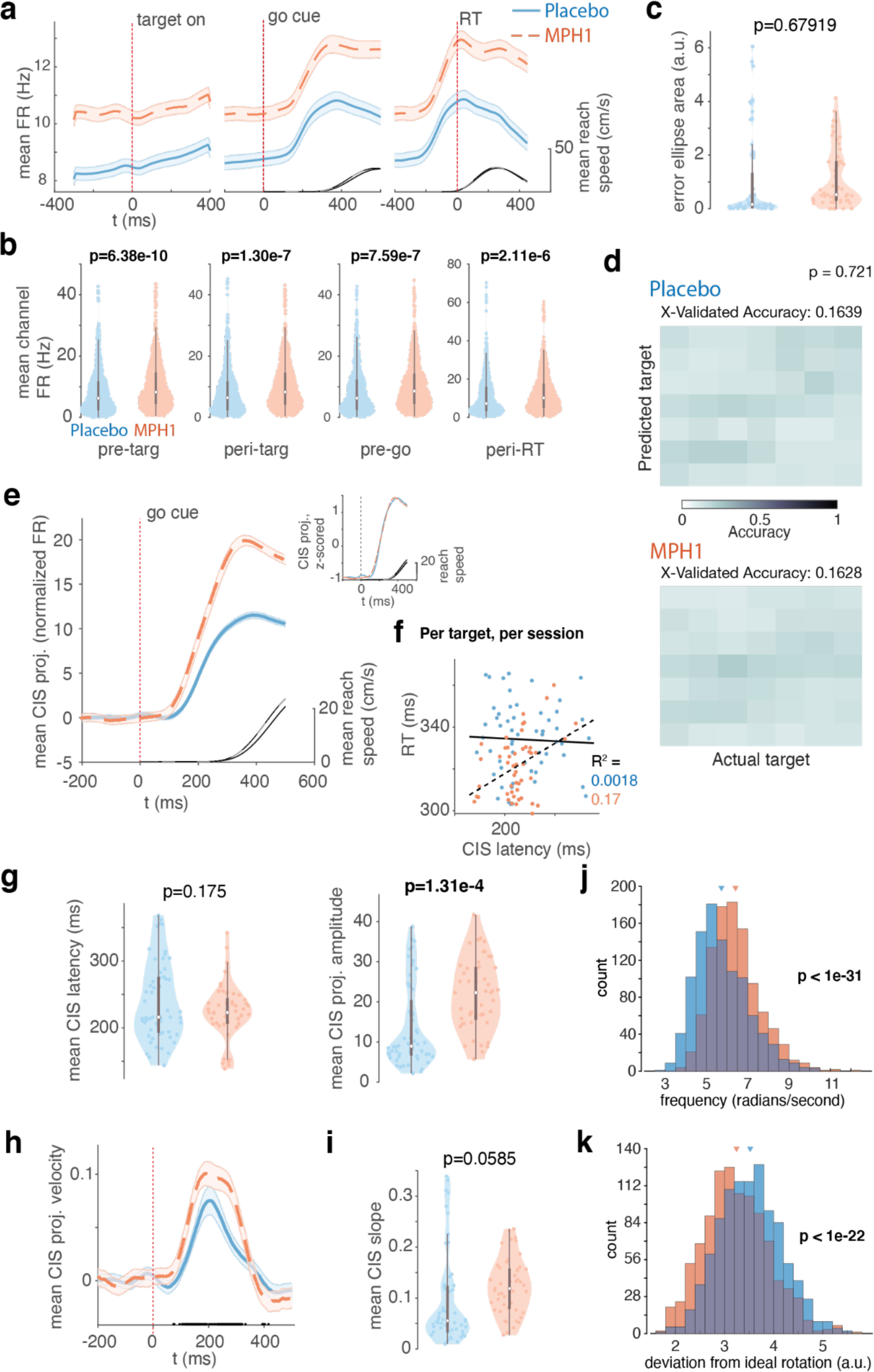
Neural effects of MPH for the P1 dataset. **a) Mean population FR trajectories by treatment.** Same as Supplementary Figure 5b, but for P1. **b) Distribution of mean single-channel PSTH values across all time points in each of four trial epochs.** Same as Figure 4d, but for P1. **c) Distributions (by treatment) of the per-session, per-target variability of the preparatory neural state.** Same as Figure 5c, but for P1. **d) Discrete reach target decoding from preparatory activity across treatment conditions.** Same as Figure 5d, but for P1. **e) Condition-invariant signal (CIS) across treatment conditions.** Same as Figure 6b, but for P1. **f) CIS correlation with reaction time (RT).** Same as Figure 6c)i, but for P1. **g) CIS latency and amplitude across treatments.** Same as Figure 6d)i, but for P1. **h) CIS velocity across treatments.** Same as Figure 6e, but for P1. **i) CIS slope across treatments.** Same as Figure 6f)i, but for P1. **j) Execution-related rotation frequency across treatments.** Same as Figure 7b, but for P1. **k) Deviation from ideal rotations across treatments.** Same as Figure 7c, but for P1.

**Supplementary Figure 7:**
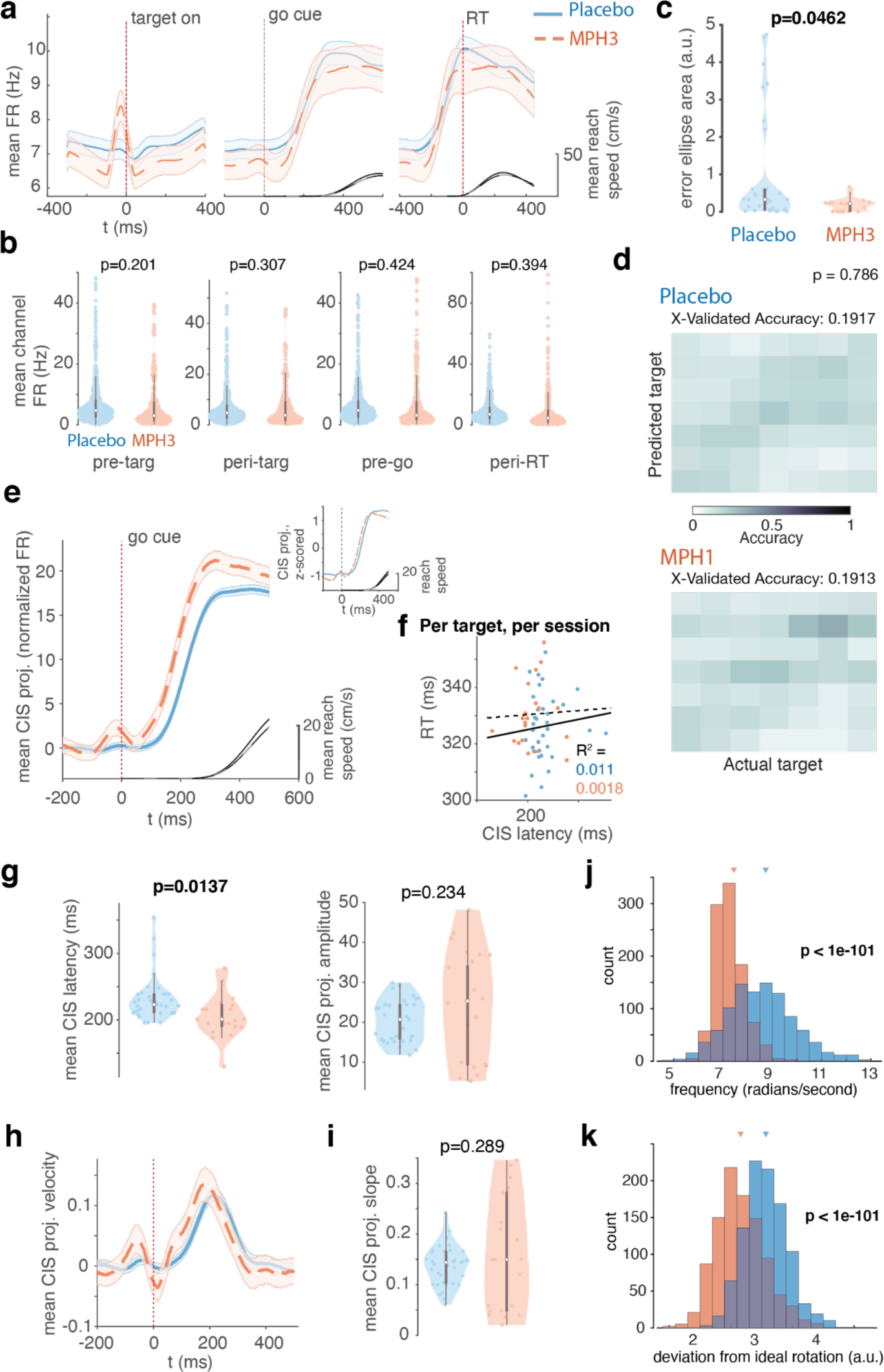
Neural effects of MPH for the P3 dataset. **a) Mean population FR trajectories by treatment.** Same as Supplementary Figure 5b, but for P3. **b) Distribution of mean single-channel PSTH values across all time points in each of four trial epochs.** Same as Figure 4d, but for P3. **c) Distributions (by treatment) of the per-session, per-target variability of the preparatory neural state.** Same as Figure 5c, but for P3. **d) Discrete reach target decoding from preparatory activity across treatment conditions.** Same as Figure 5d, but for P3. **e) Condition-invariant signal (CIS) across treatment conditions.** Same as Figure 6b, but for P3. **f) CIS correlation with reaction time (RT).** Same as Figure 6c)i, but for P3. **g) CIS latency and amplitude across treatments.** Same as Figure 6d)i, but for P3. **h) CIS velocity across treatments.** Same as Figure 6e, but for P3. **i) CIS slope across treatments.** Same as Figure 6f)i, but for P3. **j) Execution-related rotation frequency across treatments.** Same as Figure 7b, but for P3. **k) Deviation from ideal rotations across treatments.** Same as Figure 7c, but for P3.

**Supplementary Figure 8:**
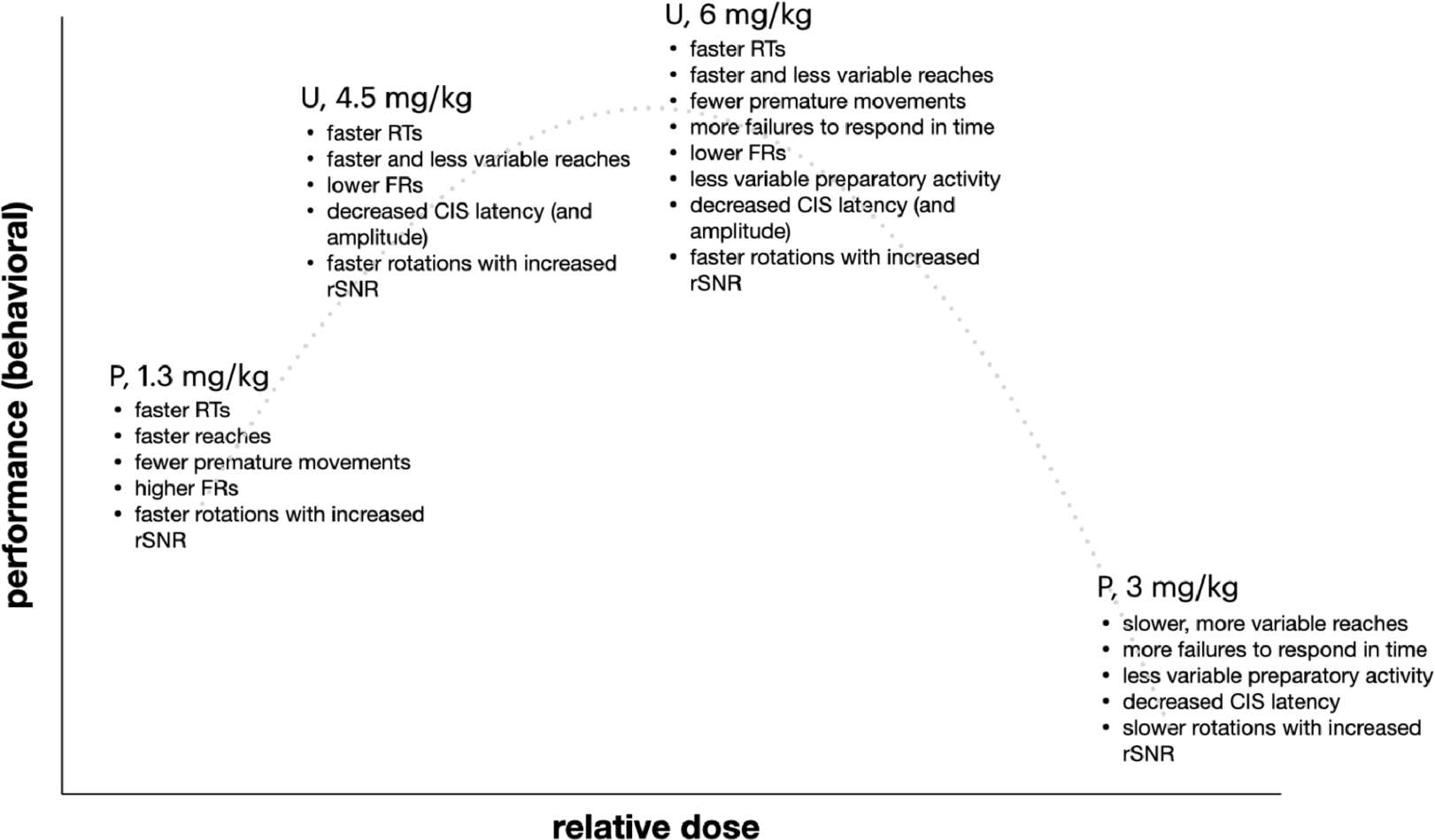
Postulated inter-subject “inverted-U” dose-response curve for effects of MPH. Summary of behavioral and neural effects by dose and monkey.

**Supplementary Figure 9:**
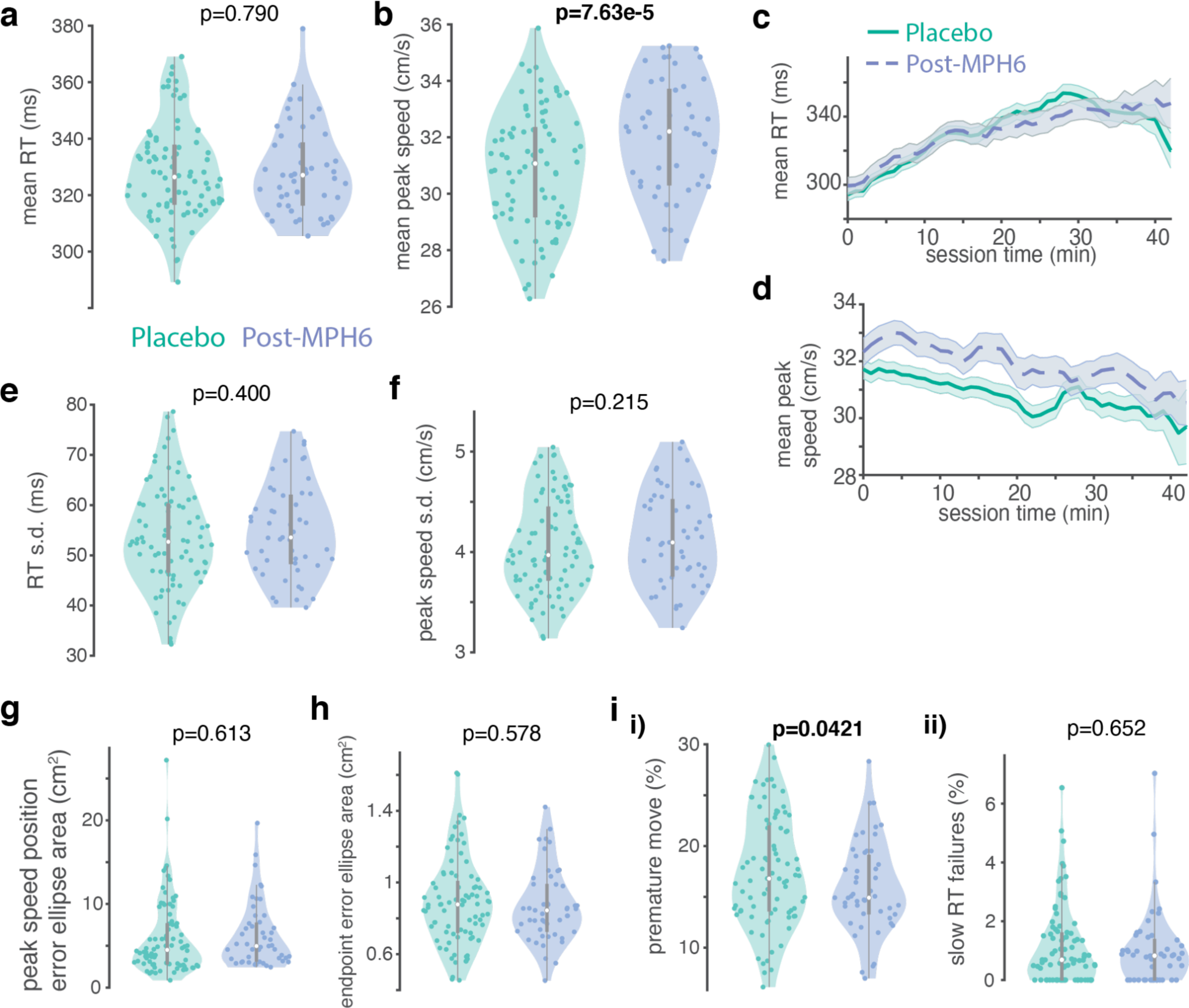
Behavioral effects of MPH in monkey U on RT, reach speed, reach variability, and trial outcomes for sessions the day after a 6 mg/kg dose and associated placebo sessions (“post-MPH6” dataset). **a) Reaction time distributions.** Same as Figure 2a, but for post-MPH6. **c) Peak reach speed distributions.** Same as Figure 2c, but for post-MPH6. **c) RT over time.** Same as Figure 2b)i, but for post-MPH. Purple dashed lines: post-MPH6 sessions; green solid lines: placebo sessions. **d) Peak speed over time.** Same as c, but for peak reach speed. **e) RT variability.** Same as Figure 3a, but for post-MPH6. **f) Peak reach speed variability.** Same as e, but for peak reach speed. **g) Variability of hand position at the time of peak reach speed across treatments.** Same as Figure 3d, but for post-MPH6. **h) Variability of reach endpoint position across treatments.** Same as Figure 3e, but for post-MPH6. **i) Impulsivity. i) False starts across treatments.** Same as Supplementary Figure 4b)i, but for post-MPH6. **ii) Sluggish starts across treatments.** Same as Supplementary Figure 4b)ii, but for post-MPH6.

